# A suite of rare microbes interacts with a dominant, heritable, fungal endophyte to influence plant trait expression

**DOI:** 10.1101/608729

**Authors:** Joshua G. Harrison, Lyra P. Beltran, C. Alex Buerkle, Daniel Cook, Dale R. Gardner, Thomas L. Parchman, Simon R. Poulson, Matthew L. Forister

**Affiliations:** Department of Botany, University of Wyoming, Laramie, WY 82071, USA; Ecology, Evolution, and Conservation Biology Program, Biology Department, University of Nevada, Reno, NV 89557, USA; Poisonous Plant Research Laboratory, Agricultural Research Service, United States Department of Agriculture, Logan, UT 84341, USA; Department of Geological Sciences & Engineering, University of Nevada, Reno, NV 89557, USA

**Keywords:** endophytes, plant traits, *Astragalus*, locoweed, swainsonine, microbial ecology

## Abstract

Endophytes are microbes that live, for at least a portion of their life history, within plant tissues. Endophyte assemblages are often composed of a few abundant taxa and many infrequently-observed, rare taxa. The ways in which most endophytes affect host phenotype are unknown; however, certain dominant endophytes can influence plants in ecologically meaningful ways–including by affecting growth and contributing to immune responses. In contrast, the effects of rare endophytes have been unexplored, and how rare and common endophytes might interact is also unknown. Here, we manipulate both the suite of rare foliar endophytes (including both fungi and bacteria) and *Alternaria fulva–a* dominant, vertically-transmitted fungus–within the fabaceous forb *Astragalus lentiginosus.* We report that rare, low-biomass endophytes affected host size and foliar %N, but only when the dominant fungal endophyte (*A. fulva)* was not present. *A. fulva* also reduced plant size and %N, but these deleterious effects on the host could be offset by a striking antagonism we observed between this heritable fungus and a foliar pathogen. These results are unusual in that they are derived from experimental manipulation in a non-crop or non-grass system and demonstrate that interactions among taxa determine the net effect of endophytic assemblages on their hosts. We suggest that the myriad infrequently-observed endophytes within plant leaves may be more than a collection of uninfluential, commensal organisms, but instead have meaningful ecological roles.

## Introduction

Plants are intimately associated with numerous fungi and bacteria that live within their tissues [1–3]. These microbes, termed endophytes [4–7] are ubiquitous and occur in hosts representing all major lineages of plants [4, 8, 9]. Over the last twenty years, it has become clear that dominant endophytic taxa can have dramatic ecological consequences–a finding demonstrated particularly well in studies manipulating the presence of vertically-transmitted fungi occurring within cool-season, perennial grasses [10, 11]. For example, these fungi can influence the successional trajectories of vegetation communities [12, 13], reshape host-associated arthropod assemblages [14, 15], and mediate host reproductive output [16]. In contrast, the ecological roles of rare endophytes–those taxa that are infrequently encountered and of low biomass–remain largely unexamined, and this is especially true of microbial associates in above-ground plant tissues. Here we manipulate assemblages of both rare and dominant endophytes living within a perennial forb to understand how these microbes interact to affect host phenotype.

Most endophytes are horizontally-transmitted among mature hosts via rainfall, air currents, or arthropods [2, 17, 18]–though facultative vertical-transmission in seeds is likely more common than often assumed [19, 20]. Many horizontally-transmitted endophytes colonize only a few cubic millimeters of host tissue [21] and may not be prevalent across samples from the same substrate [9]. Given the low biomass of these rare taxa, it is tempting to downplay their importance. However, examples from macroorganism community ecology demonstrate that certain “keystone” species, despite relatively low abundance, can exert community-wide influence [22, 23]. For instance, beavers are uncommon mammals, yet, by reshaping fluvial geomorphology, they have profound influence on co-occurring aquatic animals, waterfowl, and riparian plants [24, 25]. Similarly, rare endophytes could function as keystone species via several mechanisms, including by influencing the host phenotype, catabolism of low-concentration compounds into products required by other microbial taxa, or synthesis of potent bioactive compounds [26–28].

However, the ecological influence of rare endophytes need not be the purview of just a few species. Instead, minor effects of individual taxa could accrue to the point of assemblage-wide relevance–just as numerous genetic variants, each of minimal influence, commonly underlie phenotypes [29]. For example, an individual endophytic bacterium may trigger a highly-localized immune response of negligible importance for the host and co-occurring endophytes. But the combined effects of many bacteria can initiate systemic acquired resistance within plants, with important implications for pathogen resistance and endophyte community assembly [30–33].

Ascribing ecological influence to endophytic taxa, rare or otherwise, is complicated by a lack of understanding regarding how endophytes mediate plant trait expression [7, 34]. While the effects of certain endophytes on host growth promotion [35–37] and pathogen resistance [38–42] have attracted attention, few studies have examined endophyte mediation of other traits–including, for example, functional traits such as specific leaf area (e.g. [43]), phenology [44], and foliar elemental concentration [45] (for more see reviews by [7, 34, 46]). Nevertheless, the handful of studies demonstrating plant trait mediation by endophytes are impressive. For instance, Mejía et al. 2014 [47] report that inoculation of *Theobroma cacoa* trees with the widespread, horizontally-transmitted, fungal endophyte *Colletotrichum tropicale* affected expression of hundreds of host genes–including upregulation of some involved in the ethylene-driven immune response. These authors also found that inoculation decreased photosynthetic rate, increased leaf cellulose and lignin content and shifted foliar isotopic ratios of nitrogen (N) and carbon (C). Similarly impressive results were reported by Dupont et al.[48] who found colonization of the grass *Lollium perenne* by the *Epichlöe festucae* endophyte affected transcription of 1*/*3 of host genes (for slightly more tempered, but still dramatic, results see [49]).

Here, we manipulate the microbial consortium within the Fabaceaous forb *Astragalus lentiginosus* (spotted locoweed) to understand how endophytes belonging to different abundance categories affect plant trait expression. *A. lentiginosus* is a widespread, long-lived perennial forb that grows throughout the arid regions of the western United States of America [50]. *A. lentiginosus* exhibits extreme phenotypic variation and has over forty varietal designations [50], making it the most taxonomically rich plant species in North America [51]. The dominant fungal endophyte present within *A. lentiginosus* is *Alternaria fulva* (Ascomycota: Dothideomycetes: Pleosporaceae: *Alternaria* section *Undifilum*[52–56]). *A. fulva* is a seed-borne endophyte that grows systemically through its host and synthesizes the bioactive alkaloid swainsonine [57]. Consumption of swainsonine-laced tissues by mammalian herbivores can lead to extreme toxicosis and even death [58]. *A. fulva* is prevalent throughout the range of its host and thus could be considered a “core” taxon within this substrate [59], though not all populations of *A. lentiginosus* are colonized by the fungus, and intrapopulation variation in fungal colonization has also been reported [56, 60–63].

*Alternaria* section *Undifilum* fungi have been observed in numerous swainsonine-containing taxa within *Astragalus* and *Oxytropis* that are colloquially called “locoweeds” [54, 64–66]. The nature of the relationship between locoweeds and their seed-borne fungi is somewhat unclear. Swainsonine production does not seem to influence certain specialist arthropod herbivores [67, 68] and may, in some cases, increase mammalian herbivory [69]. These observations have led Creamer et al.[70] to hypothesize that *Alternaria* section *Undifilum* taxa live as commensals within locoweeds. However, recent work supports a more mutualistic relationship. For instance, Harrison et al.[56] demonstrated, via a DNA sequence-based survey, that swainsonine concentrations and *A. fulva* relative abundance were inversely related to fungal endophyte richness, potentially reducing exposure of hosts to pathogens. In a culture-based survey, Lu et al. [55] reported similar results in two other locoweed species (also see [71] for an analogous phenomenon in a *grass-Neotyphodium* endophyte system). The results from these surveys suggest that vertically-transmitted *Alternaria* endophytes can shape fungal endophyte assemblages, though effects on bacterial endophytes are unknown. Additionally, Cook et al. 2017 [72] demonstrated that *Alternaria* section *Undifilum* endophytes can affect the biomass and protein content of several locoweed taxa, including *A. lentiginosus.* These results suggest *A. fulva* may mediate other host traits as well.

By removing embroyos from the seed coat, the abundance of *A. fulva* in plant tissues can be greatly reduced [73]. We used this approach to manipulate the presence of *A. fulva* in *A. lentiginosus* plants to experimentally test the aforementioned antagonistic relationship between *A. fulva* and co-occurring endophytes and explore how *A. fulva* affects various host traits, including size, leaflet area, specific leaf area, foliar C and N, phenology, and rhizosphere activity. For a subset of focal plants, we applied an inoculum slurry to leaf surfaces to increase overall endophyte diversity and boost the occurrence of rare, horizontally-transmitted taxa. We applied these manipulations in a full factorial design to compare how endophytes of differing abundance categories shape host plant phenotype, and explore how a dominant endophyte might influence co-occurring fungi and bacteria.

## Methods

### Field experiment

During the early spring of 2017, seeds of *A. lentiginosus* var. *wahweapensis* from the Henry Mts. UT, U.S.A. (collected in 2005 from a population known to possess *A. fulva*) were lightly scarified, left to imbibe deionized water overnight and germinate indoors in a mix of humus, compost, and topsoil sourced from the Reno, NV region. To reduce the relative abundance of the vertically-transmitted fungal endophyte, *A. fulva*, embryos were excised from a subset of seeds prior to planting, as per [73]. Seedlings were grown at ambient temperature under a 16:8 (light:dark) lighting regime and watered with dechlorinated tapwater. Individuals from different treatments were interspersed haphazardly, but not allowed to touch one another. Seedlings were periodically reorganized to avoid any influence of subtly differing conditions across the growth area. To speed growth, Miracle-Gro (Scotts Miracle-Gro Company, Marysville, OH) was applied several times to all replicates during the first month of growth. To control for possible confounding effects of seed coat removal, embryos were excised from a subset of seeds and planted along with potato dextrose agar (PDA) that was sterile, or that was colonized by *A. fulva*, which had been cultured from intact seeds. These control seedlings were planted several weeks later than other seedlings, due to slow growth of *A. fulva* cultures.

In early June, seedlings were installed in five gallon pots filled with equal parts locally sourced humus and topsoil and placed in an abandoned, largely denuded, field near the University of Nevada, Reno. A total of 300 plants were installed (between 54–68 per treatment group, see Table S1). Pots were organized randomly with respect to treatment and were placed one meter apart so plants never touched one another. Dechlorinated water was applied as needed to all plants at the same time (typically every other day, except during the heat of summer when watering was conducted daily). Every two weeks a slurry of microbial inocula (described below) was sprayed on leaves of half of the plants. A solution with identical surfactant, but no microbial inoculum, was applied to untreated plants. Plants were left in the field from early June through mid-September, at which point leaves were removed for sequencing and culturing.

### Inoculum preparation

Foliar microbes were isolated from the following woody shrubs growing near Reno, NV: *Artemisia tridentata*, *Ericameria nauseosa*, *Prunus andersonii*, and *Tetradymia canescens*. These shrubs are abundant throughout the Great Basin Desert and, consequently, we reasoned they contained horizontally-transmitted, foliar microbes likely to be regularly encountered by *A. lentiginosus*. Individual shrubs to be sampled were selected haphazardly. We did not use leaves from *Astragalus* species to avoid inoculating plants with *A. fulva* and thus obviating our treatment to reduce this fungus. Leaves were cut into sections (of several mm^2^) and placed on PDA and the resulting microbial growth isolated and subcultured. Twenty morphologically unique, reproductive isolates were used to make inoculum, including 11 isolates from *A. tridentata.* Spores were removed and suspended in deionized water and 0.0001% TWEEN 20®(Sigma-Aldrich), a detergent that functioned as a surfactant. A haemocytometer was used to dilute the suspension to ~100,000 spores ml^-1^. This concentration was chosen because it produced no obvious negative symptoms in *A. lentiginosus* seedlings during preliminary experiments. Aseptic technique was used throughout culturing and inoculum preparation. Two aliqouts of inoculum were sequenced to identify the constituent microbial taxa (DNA sequencing methods for characterizing endophyte communities are described below).

### Plant trait measurement

All plant traits were measured concomitant with sample collection for foliar microbiome characterization. Plant size was measured as the product of the width at widest point, the width perpendicular to that point, and plant height. Phenological state and number of leaves were characterized for each plant. Area and specific leaf area (SLA; leaflet area divided by mass of leaflet) were measured for three dried leaflets per plant and averaged. Two healthy leaflets were removed from 8–12 leaves per plant, rinsed with tap water, dried in a laminar flow hood (<12 h total) and frozen until further processing. These leaflets were then parsed for microbiome characterization, swainsonine quantification, and carbon (C) and nitrogen (N) analysis. Swainsonine concentration in ~50 mg of dried, ground foliar tissue was measured using an LC-MS/MS approach described by [60]. Briefly, an 18 h extraction in 2% acetic acid with agitation was followed by centrifugation. Supernatant was added to 20 mM ammonium acetate and subjected to LC-MS/MS analysis. Percent C and N and ^15^N:^14^N isotopic ratios in 3–4 mg dried foliar tissue, were measured by the Nevada Stable Isotope Laboratory using a Micromass Isoprime stable isotope ratio mass spectrometer (Elementar, Stockport, UK) and a Eurovector elemental analyzer (Eurovector, Pavia, Italy). The percentage of nitrogen in tissues due to fixation alone (NDFA) was calculated as per [74] through comparison with samples from co-occurring *Chenopodium album,* which is not known to harbor nitrogen fixing rhizosphere bacteria (for further details see [75]).

### Sequence and culture-based characterization of the foliar microbiome

We characterized endophytic assemblages through both culturing and DNA sequencing, thus affording us insight into the effects of treatment on the microbiome from two independent techniques. For our culture-based assay, we choose three leaflets per plant. Leaflets were surface sterilized, cut into 3–4 pieces, and plated onto PDA, using aseptic technique. Surface sterilization involved rinsing in 95% ethanol for 30 s, 2 min in 10% sodium hypochlorite solution, 2 min in 70% in ethanol, followed by a final rinse in deionized water. Preliminary experiments confirmed the success of this surface sterilization technique (data not shown). Cultures were grown in the dark at ambient temperatures for 1.5 months. Microbial growth was isolated, subcultured, and the number of morphologically unique cultures and the percentage of leaf pieces colonized recorded. Cultures corresponding to *A. fulva* were identified visually through comparison to *A. fulva* cultures grown from seeds used for this experiment. Culture identification was confirmed via sequencing.

DNA was extracted from three surface-sterilized, dried, and ground leaflets per plant using DNeasy plant mini kits (Qiagen, Hilden, Germany). An extraction blank for each kit was generated and blanks used as negative controls. Library preparation and 2×250 paired- end sequencing on the Illumina MiSeq platform (San Diego, CA, U.S.A.) was performed by the Genome Sequencing and Analysis Facility at the University of Texas, Austin, U.S.A. To characterize bacterial assemblages, the 16s (V4) locus was amplified using the 515-806 primer pair [76], while for fungal assemblages the ITS1 locus was amplified using the ITS1f- ITS2 primer pair [77]. PCR reactions were performed in triplicate, with both positive and negative controls. A mock community consisting of eight bacteria and two fungi was also sequenced (Zymo Research, Irvine, CA. U.S.A.) as an additional positive control and a way to determine suitability of our bioinformatic approach. Extracted DNA from five plants was parsed into technical replicates, passed through library preparation, and sequenced to determine variation due to instrumentation. For full library preparation details see the supplemental material.

Sequences were merged with USEARCH v10.0.240 [78] using default parameters. Primer binding regions were stripped from merged reads, as these regions are more likely to contain sequencing errors. The expected number of errors within each read was estimated based on quality scores and reads with more than a single expected error discarded [79]. Unique reads were clustered into exact sequence variants (ESVs) using UNOISE3 [80]. ESVs offer numerous advantages to 97% OTUs [81] (for further discussion see the Supplemental Methods). ESVs that were shorter than 64 bases long, consisting of repeated motifs, or that corresponded with PhiX were removed from the data. Unfiltered reads were matched to consensus ESV sequences and an ESV table generated via USEARCH, thus salvaging some data from reads that did not pass our stringent filtering. If *>* 1% of the total reads for an ESV were in the negative control, then these ESVs were deemed possible contaminants and discarded.

Taxonomic hypotheses were generated for ESVs using the SINTAX algorithm within the USEARCH software [82]. For ITS data, the UNITE (v7.2; [83]) and Warcup (v2; [84]) training databases were used, and for 16s data the Greengenes (v13.5; [85]) and Ribosomal Database Project (v16; [86]) training databases were used. For each marker, taxonomic hypotheses generated by training databases were merged and the lowest level of taxonomic agreement identified. To remove host DNA from 16s data, ESVs were queried against the MIDORI [87], MITOFUN [88], and RefSeq mitochondria and plastid databases [89] and ESVs matching mtDNA or cpDNA within any database were removed. For ITS data, ESVs identified as plant ITS via comparison to the UNITE and Warcup databases were removed. ESVs corresponding to *A. fulva* were identified through comparison to GenBank accession JX827264.1.

### Statistical analysis

We analyzed data output by our sequencing approach via a hierarchical Bayesian modeling (HBM) framework that provides estimates of proportional relative abundance for each microbial taxon (following [90]). The model estimates parameters of replicate-specific, multinomial distributions that describe taxon proportions *(p* parameters) and Dirichlet parameters that describe proportion estimates for the entire sampling group ([90]). This method shares information among replicates for more accurate parameter estimation and allows propagation of uncertainty in parameter estimates to downstream analyses. Modeling was conducted in the R computing environment (R Core Team 2017) using the JAGS model specification language [92] as implemented via rjags v4-6 ([93]; for full model description see code provided in the Electronic Supplementary Material). The Markov chain Monte Carlo (MCMC) sampler in rjags was adapted until it approached the theoretical optimum or for 30,000 iterations, whichever came first. The first 100,000 MCMC samples of the two chain model were discarded (burn in period) and every 4th sample of the next 4000 samples retained. Modeling generated MCMC samples that characterized the posterior probability distributions (PPDs) for focal parameters. MCMC convergence was tested using the Gelman-Rubin and Geweke statistics [94, 95]. Models were implemented with and without *A. fulva* sequences. The latter approach was necessary to compare the relative abundances of non-*A. fulva* fungi among treatment groups, because the abundance of *A. fulva*, when it was present, shifted the relative abundance of all co-occurring taxa, which could cause spurious results [96]. To determine if ESVs differed in relative abundance among treatment groups, the degree of overlap of PPDs for Dirichlet parameters for each ESV was calculated. If PPDs overlapped by 95% or more then there was little evidence for an effect of treatment, if, however, PPDs barely overlapped, then a shift in microbial relative abundance associated with treatment was credible.

Species equivalents of Shannon’s entropy and Simpson’s diversity [97] were calculated at two levels–for the treatment group as a whole and for each replicate. To estimate diversity equivalencies for a treatment group, the equivalency was calculated for *each* MCMC sample of the Dirichlet distribution characterizing microbial relative abundances within that treatment group. This generated a PPD of diversity, thus propagating uncertainty in relative abundance estimates into estimates of treatment group diversity (for a similar approach see [98]). To determine to what extent diversity equivalents differed between treatment groups, the overlap of PPDs for each group was examined (as per above). Diversity equivalents were also estimated for each replicate so that these estimates could function as the response in a linear model testing for associations between plant trait variation and shifts in microbial diversity (see below). To estimate diversity for each replicate, the means of PPDs of multinomial parameters for that replicate were calculated (recall that these parameters estimated proportional microbial relative abundance). Diversity equivalencies were calculated for the resulting vector. ESVs corresponding to *A. fulva* were not included in calculations at either the replicate or treatment group level.

HBM was also used to estimate differences among treatment groups in the mean values of plant traits, sequence-based estimates of microbial diversity, and culture richness. Each response variable was modeled as a draw from a normal distribution characteristic of the sampling group as per [99]. The mean *(μ)* and variance *(τ*) of this distribution was estimated through sharing of information among replicates. The prior distribution for *μ* was a normal distribution centered at zero with a precision of 0.0001. The prior distribution for *τ* was a uniform distribution from 0–100 (for full model specification see R code provided in the Electronic Supplemental Material). MCMC sampling and tests for credible effects of treatment via PPD overlap were conducted as described above.

To evaluate associations between plant traits and microbial diversity, linear models were created in a HBM framework. Beta coefficients for plant traits were estimated for each treatment group, with a prior sampled from a normal distribution centered at the estimated across-treatment effect of each trait and a precision estimated across all treatments. Hyperpriors for beta coefficients were normal distributions centered at zero with a precision of 0.0001 (for full model specification see R code provided in the Electronic Supplemental Material; also see [56]). Means of PPDs for each beta coefficient were used as point estimates of the effect of that covariate. The proportion of the PPD for each beta coefficient that did not overlap zero was used to determine certainty of a non-zero effect. Prior to modeling, missing values in covariates were imputed using the random forest algorithm [100] as implemented by the randomForest R package [101]. When models were run without imputing data, results were similar to those reported here.

We refrain from presenting taxon-specific analyses here that would link the relative abundance of specific endophytes to plant traits for two reasons. First, endophytes were manipulated as a suite. Consequently, all members of the suite will be associated with shifts in host phenotype due to treatment, increasing the chances of spurious correlation and making keystone identification more challenging. Second, the compositional nature of sequencing data renders taxon-specific analyses fraught, particularly when the microbial community is dominated by one species, which is itself subject to manipulation [96, 102, 103]. For the same reasons, we chose not to report effects of treatment on endophyte richness. To explain, when *A. fulva* is present it is typically dominant and many of the fungal reads are assigned to this taxon. Therefore, even if *A. fulva* is removed from the data, samples from plants containing the taxon would have fewer reads available to allocate to the other microbes present, thus reducing richness artificially compared to plants without *A. fulva.* To assay effects of *A. fulva* presence on richness, we therefore rely on data obtained through culturing.

We omitted from all analyses those few plants for which seed coat removal did not remove *A. fulva* as ascertained via swainsonine concentration (this compound is not known to be produced by the host plant). We also omitted those plants with no evidence of *A. fulva* occurrence from the *A. fulva* positive treatment group because *A. fulva* is known to be incompletely transmitted between generations [62, 104, 105], thus some seeds were expected to be uncolonized or to possess only trace amounts of the fungus. Analyses were repeated while retaining all of these plants and the results obtained were qualitatively similar to those presented here.

## Results

### Effects of the vertically-transmitted fungus on the host and co-occurring microbes

Treatment to reduce the relative abundance of the dominant, heritable fungus *A. fulva* from *A. lentiginosus* plants was successful as evidenced by read counts (Fig. S1), swainsonine concentrations (Fig. 1), and culturing efforts (Fig. S2). *A. fulva* presence influenced plant phenotype–colonized plants were much smaller and had fewer leaves than uncolonized plants (Figs. 1; S4). The negative effect of *A. fulva* on plant size was observed in the second year of monitoring as well (Fig. S3). Foliar N was affected by both *A. fulva* and rare microbes–plants without *A. fulva* and that were untreated with inoculum had elevated %N in their leaves. Moreover, *A. fulva* increased the ^15^*Ĵ* (ratio of N isotopes) and reduced NDFA,a proxy for rhizosphere activity. The effects of *A. fulva* colonization on host phenotype were apparent in control treatments, thus the observed results are not due to the confounding influence of seed coat removal (Fig. S4). We did not observe an effect of *A. fulva* colonization on %C or SLA.

**Figure 1:**
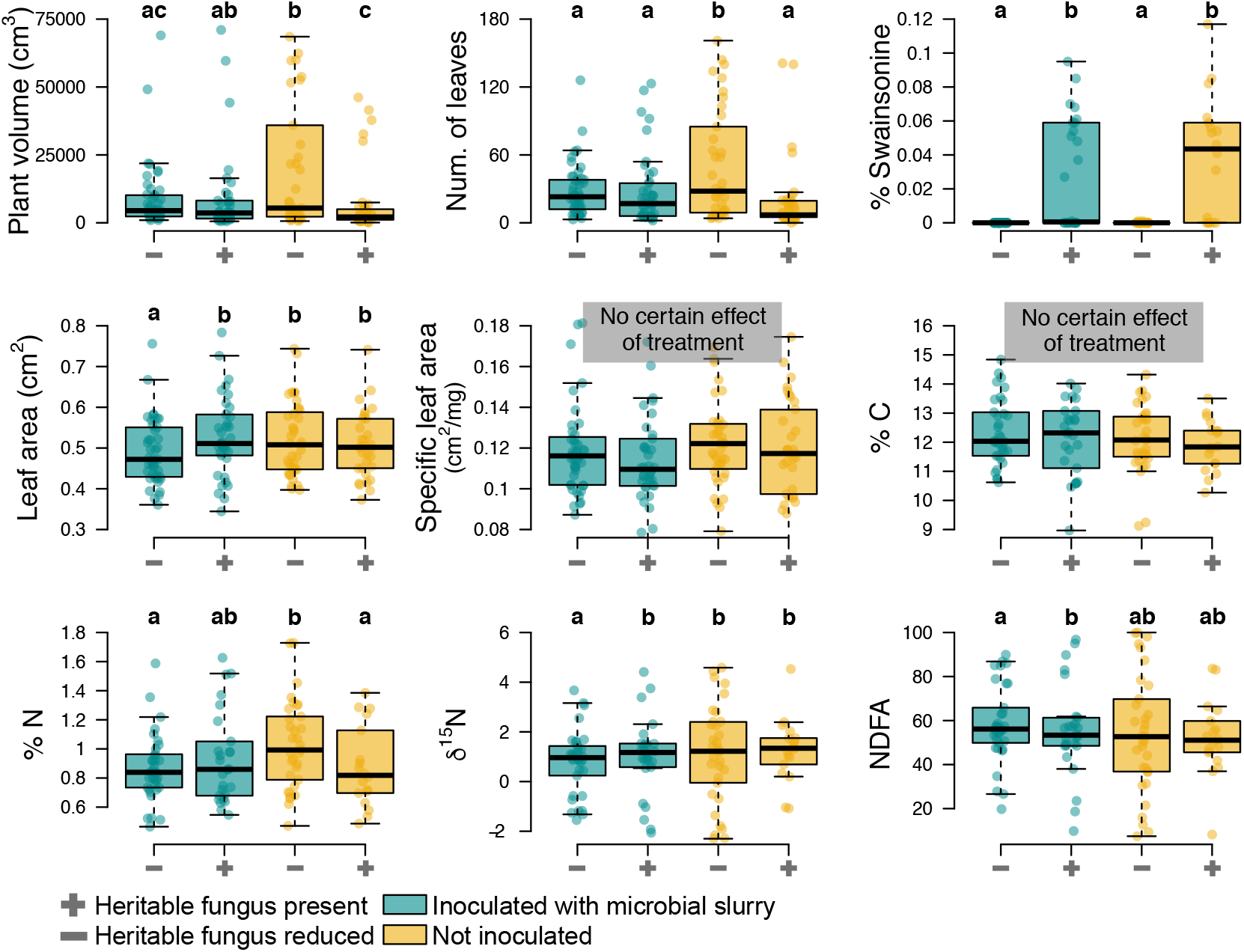
Variation in *A. lentiginosus* traits among treatment groups. + and - symbols on the x-axis denote treatment to reduce the relative abundance of the vertically-transmitted fungus, *A. fulva*. Boxes shaded blue denote treatment with endophyte inoculum slurry; boxes shaded yellow denote plants that did not receive the slurry. Percentage of N, C, and swainsonine refer to foliar dry mass composition. Differences in mean trait values among treatment groups were determined through a hierarchical Bayesian analysis. Credible differences among treatment groups are denoted through the letters above each boxplot. For estimates of mean trait values for each treatment group see Table S2. Boxplots summarize the data and describe interquartile range with a horizontal line denoting the median. Whiskers extend to the 10th and 90th percentiles. Several outliers were omitted to aid visualization.

*A. fulva* presence had little effect on co-occurring bacteria, but strongly influenced other fungal endophytes (Table S3, Figs. 2, S5). Indeed, in most cases, when we observed fungi differing in relative abundance among treatment groups, those groups also differed in *A. fulva* colonization (Table. S3). Notably, we conducted analyses while omitting *A. fulva* so these results are unlikely to be artifacts caused by the compositional nature of sequencing data [96]. *A. fulva* presence also influenced overall fungal diversity. Species equivalents of Shannon’s entropy increased in plants colonized by *A. fulva* (Fig. 2), but the opposite was true for equivalents of Simpson’s diversity. Simpson’s diversity index places more weight on abundant taxa than does the Shannon index [106]. These contrasting results appear to be driven by a negative association between *A. fulva* and the dominant ESV assigned to *Leveillula taurica. L. taurica* is a powdery mildew (Erysiphaceae) known to colonize numerous plant species, including certain *Astragalus* taxa [107, 108]. *L. taurica* was the second most abundant fungal endophyte sequenced (behind *A. fulva*) and dropped in relative abundance when *A. fulva* was present (Fig. S7). This antagonism was also observed visually, as we noted a powdery mildew infection on the leaves of a subset of the plants used for this experiment, and infections were less severe in plants colonized by *A. fulva* (Fig. S6). For less abundant fungal endophytes, including uncommon genotypes of *L. taurica*, we observed an increase in relative abundance when *A. fulva* was present (Fig. S7), which is likely responsible for the increase in Shannon’s diversity with *A. fulva* infection. Data from culturing concurred with those obtained from sequencing–fungal endophyte richness and colonization tended to increase with *A. fulva* colonization (Fig. S2).

**Figure 2:**
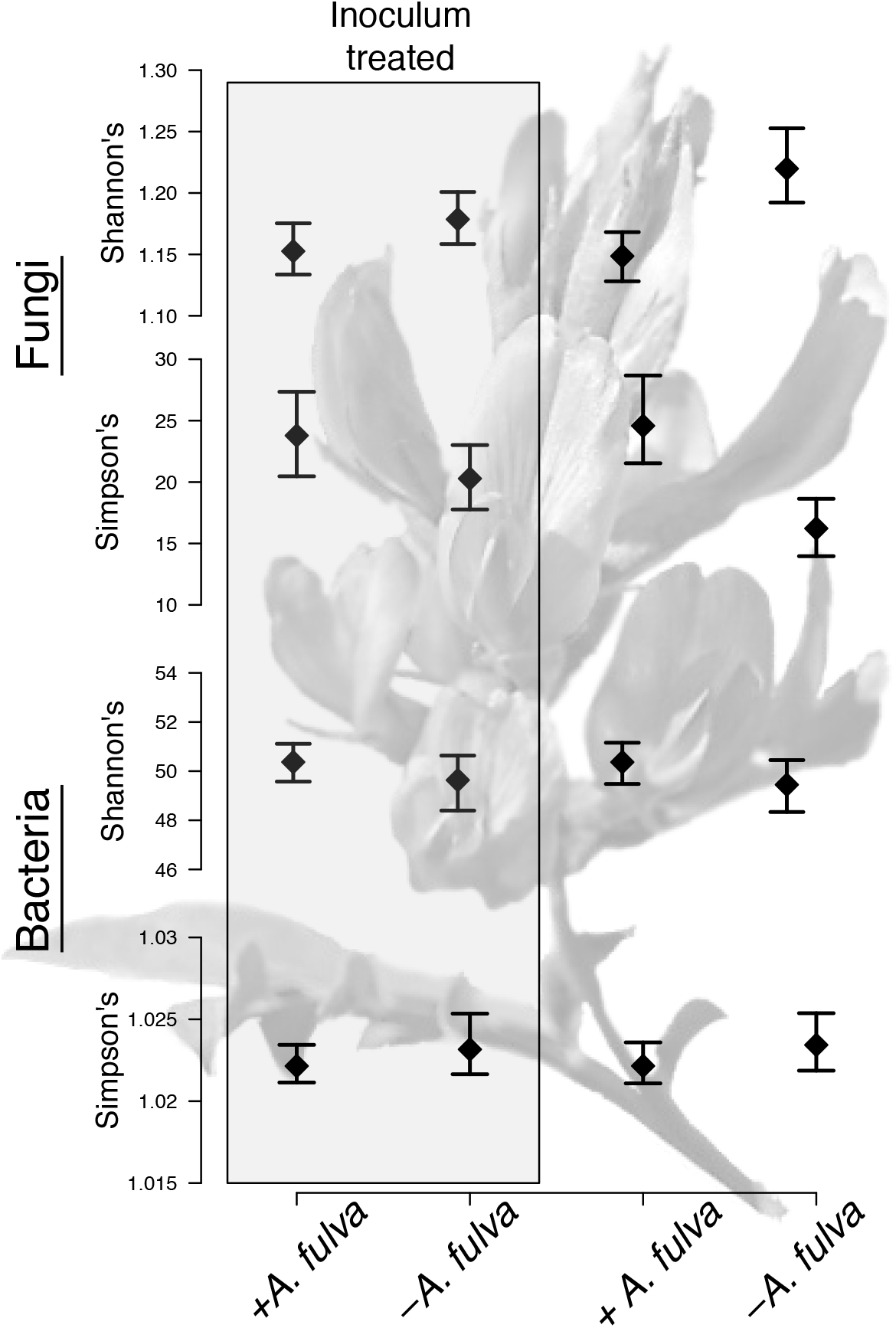
Influence of treatment on fungal and bacterial diversity. Estimates of species equivalencies of Shannon’s entropy and Simpson’s diversity are shown for each treatment group. Inoculum treated groups are enclosed in the gray box. Points denote the means of posterior probability distributions (PPD) of diversity equivalencies. Line segments extending from means denote the 5–95% credible intervals of the PPD. A rendering of *A. lentiginosus* underlies the plot (photo by J. Harrison).

### The influence of horizontally-transmitted endophytes on hosts

The inoculum slurry applied to plants was created from twenty morphologically distinct cultures derived from shrubs common in the Great Basin Desert (see methods). Sequencing revealed the slurry was composed of ten fungal and 88 bacterial ESVs–36 of these bacterial ESVs and four of the fungal ESVs were sequenced from plants used in this experiment. In almost all cases, inoculum application led to an increase in the read counts obtained for taxa within the inoculum mixture (in 100% of fungal taxa and 86% of bacterial taxa), suggesting treatment was successful. Additional support for the efficacy of inoculation was obtained from our culturing efforts–we observed increased culture morphospecies richness for plants treated with inoculum than for untreated plants (Fig. S2). Fungi present in inoculum that were also present in treated plants were from the Dothideomycetes, while successful bacterial colonizers were predominantly members of Proteobacteria (15 ESVs), Actinobacteria (8 ESVs), and Firmicutes (8 ESVs). Inoculum application had no effect on overall microbial diversity (Fig. 2).

Inoculum application had no visibly pathogenic effects on plants–they appeared healthy and leaves had no evidence of necrosis. However, inoculum application did influence plant phenotype, but only when the dominant fungus *A. fulva* was not present. For instance, inoculum application reduced plant biomass, foliar %N, and caused plants to have approximately 50% fewer, smaller leaves (Table S2), but this was only apparent for plants without *A. fulva* (Fig. 1). In general, associations between the diversity of horizontally-transmitted endophytes, either bacterial or fungal, with plant trait variation were weak and often limited to a specific treatment group (Tables S4, S5).

### Sequencing summary and microbial diversity description

After removing host reads and applying our stringent quality control approach, we retained for analysis 115,860 reads from 11 fungal ESVs and 1,496 reads from 53 bacterial ESVs (for full details see the Supplemental Results). The low number of bacterial reads reflected a dominance of 16s reads by host chloroplast DNA and was not due to faulty library preparation efforts (see Supplemental Results). The proportion of reads assigned to an ESV varied less than <1% among technical replicates, and patterns of ESV occurrence among technical replicates were nearly identical. Additionally, our bioinformatics approach did well capturing diversity present in the mock community (see Supplemental Methods).

The majority of fungal ESVs belonged to the Ascomycota, with the Dothideomycetes and Leotiomycetes particularly well represented (4 ESVs each). The Proteobacteria (18 ESVs) constituted the bulk of the bacterial taxa observed, but Firmicutes and Actinobacteria were also common (9 ESVs each). Two ESVs matched *A. fulva*, with one genotype being much more abundant than the other (Fig. S1).

## Discussion

Foliar endophyte assemblages are typically composed of few dominant, often widespread, generalist taxa (often referred to as the “core” microbiome; [59]) and numerous low-abundance taxa that occupy a minuscule proportion of their host’s tissues [21, 109, 110]. Relevance is often presumed the domain of dominant and core microbial taxa, because of their greater biomass and prevalence. Our results suggest the truth is more complex; we report that a suite of rare endophytes affected several host plant traits, including size and foliar N content. We also observed that the influence of rare endophytes was attenuated by the presence of a dominant fungal endophyte, which also mediated host plant phenotype.

It is important to note that, regardless of treatment, host plants appeared healthy to the eye–their tissues were green, and, in many cases, they fruited successfully during the first and second years of growth. Thus, both *A. fulva* and co-occurring microbes meet the criterion of living asymptomatically within plant tissues necessary for designation as endophytic taxa rather than obligate pathogens [5]. Many of the microbes constituting our inoculum mixture successfully colonized host plants, but were quite rare, as evidenced by read counts (by comparison, many more reads were obtained from *A. fulva* and *L. taurica*). Therefore, it seems likely that these taxa occupied only small portions of host tissues. Despite their low biomass, inoculation with rare microbes decreased host size, foliar N%, leaflet area, and specific leaf area (Fig. 1), but only in plants that were not colonized by *A. fulva.*

Our results compliment recent research by Christian et al.[45] who reported endophytes, and interactions between endophytes and pathogens, can alter N distribution and uptake in plants. This study did not report endophyte-induced shifts in %N content at the whole plant level, but did find endophytes increased ^15^N in plants and affected ^15^N distribution among leaves (also see [47]). We also observed subtle shifts in *Ĵ*^15^N and NDFA with treatment (Fig. 1; Table S2). NDFA is derived from ^15^N and ^14^N isotope ratios (see methods) and is a proxy for rhizosphere activity. Additionally, we report that rare endophytes reduced foliar %N, but only when *A. fulva* was not present. When taken together with previous work [45, 47, 111], these results suggest that the effects of horizontally-transmitted endophytes on foliar N depend on host, abiotic context (e.g. N availability), and interactions with other microbiota.

The effects of rare endophytes on trait expression were somewhat subtle, with the exception of striking effects on plant size, including an approximately 50% reduction in plant size and leaf number for plants without *A. fulva* (Table S2). These effects likely have implications for host fitness–in particular, a reduction in host size corresponds to reduced seed output (see below). Moreover, small changes in leaf traits caused by endophytes, including shifts in N concentration, could scale up to have considerable importance for ecosystem productivity and elemental cycling [112]. Additionally, shifts in host phenotype could have indirect effects on host-associated organisms, such as arthropods. For instance, size and foliar %N are often strong predictors of variation in insect assemblages and herbivory across plant species [75, 113, 114], thus endophyte-mediated shifts in these traits could have cascading effects on arthropod communities. As a caveat, it is plausible that our inoculum mixture was biased towards endophytes that grow rapidly in culture and slower-growing taxa could have different effects on the host than those we observed here.

We considered two possibilities for the distribution of ecological influence among rare endophytic taxa. Specifically, influence could be limited to several keystone taxa or could be cumulative, such that a quorom must be reached before the combined effect of rare endophytes induces a response by the host. We were unable to satisfactorily resolve these two non-mutually exclusive hypotheses. However, the quorom hypothesis should lead to a negative association between microbial diversity and plant size, which we did not observe (Tables S4, S5). Indeed, we found that most rare endophytes occurred infrequently in samples, suggesting that if a quorom was present and responsible for the shifts in host phenotype observed, then that quorom must be easily met and be composed of very little total biomass. Alternatively, infrequently-observed, keystone taxa could have caused the treatment effects we report, as these taxa, by definition, exert greater influence than would be predicted from their low biomass. For instance, a localized infection by a keystone taxon could have effects that spread throughout the host, yet that taxon would not be present in the majority of leaves sequenced from that host. This concept suggests limitations of the common practice of *in silico* identification of keystone taxa as those taxa that are prevalent among samples, such that their removal from co-occurrence networks causes a shift in network topology [115–117]. We reiterate the non-exclusivity of the keystone and quorom hypotheses and suggest disentangling the two represents a profitable line of inquiry for future work.

The dominant, vertically-transmitted fungal endophyte *A. fulva* also reduced host size and foliar %N (Fig. 1; consistent with [72]). Inhibition of host growth by *A. fulva* is perplexing, because the fungus is vertically-transmitted in seeds and, therefore, its fitness is tied to that of its host. Larger *A. lentiginosus* plants generally produce more seeds (J. Harrison, personal observation) and, thus, selection should operate against mechanisms by which *A. fulva* reduces host growth. On the other hand, *A. fulva* grows very slowly in culture [118, 119] and fast-growing plants could possibly outpace hyphal growth. If the fungus cannot grow fast enough to reach seeds before they mature, then its direct fitness is zero. Consequently, constraining host growth may improve fungal fitness, because it would allow time for hyphae to reach reproductive structures. This hypothesis awaits further testing. Another intriguing possibility is that a fungal-induced reduction in plant size could actually improve plant longevity in extreme conditions. For several native plants in the Great Basin, small stature paradoxically facilities the ability to withstand drought and competition from invasive annual grasses [120, 121]. Thus, it is possible that plants colonized by *Alternaria* endophytes could better survive the harsh desert climate. Interestingly, previous work has shown that *Alternaria* endophytes do not reduce plant size in locoweeds that are drought-stressed [122] and that swainsonine concentration can increase during drought stress.

The potential fitness costs imposed by *A. fulva* on its host may be further ameliorated by the obvious antagonism we observed between *A. fulva* and the most abundant pathogen present, *Leveillula taurica* (Figs. S6, S7). These results support the hypothesis posed by Lu et al.[55] that the *Alternaria* spp. occurring within *Astragalus* and *Oxytropis* act as mutualists by restricting pathogen exposure. *A. lentiginosus* is a plant of frequently disturbed, climatically variable, arid landscapes and it is likely that pathogen pressure in such locales is particularly damaging, because the lack of resources could impede recovery from tissue loss. The same rationale has inspired the growth-rate hypothesis in the plant-insect literature [123, 124]. This hypothesis predicts plants growing in resource poor conditions will recuperate from herbivory slowly, and thus benefit from investment in phytochemical defenses that are otherwise too costly. Similarly, tolls imposed by *A. fulva* on *A. lentiginosus* may be acceptable to the host given the harshness of the arid American West. Indeed, it seems likely the benefits of *A. fulva* presence outweigh the costs, because numerous populations of *A. lentiginosus* harbor the fungus [56, 61, 63].

Our study demonstrates the ecological consequences of interactions among microbes [28], as evidenced by a negligible effect of inoculum application for plants colonized by *A. fulva* and the antagonism between *A. fulva* and *L. taurica*. We suggest that these results are not likely due to direct interactions between *A. fulva* and other microbes–the disparity in leaf size and microbe size is too great. Even for *A. fulva*, which grows systemically through its host, physical encounters with co-ocurring microbes are probably rare [125, 126]–with the possible exception of encounters with *L. taurica*. Instead, we suggest microbe-microbe interactions are likely to be indirectly mediated by the host. The mechanism behind these indirect interactions is unknown. However, gene expression studies in several perennial grasses [48, 49, 127] and in *Theobroma cacao* [47] have demonstrated an upregulation in the host immune response post-colonization by endophytes (see [128]). To speculate, it is possible that *A. fulva* similarly primes the host immune response, which could negatively affect co-occurring microbes. Since fungi are often attacked by a different component of the plant immune response than bacteria [129, 130], it is likely that immune system priming by *A. fulva* would have stronger effects on other fungi than on bacteria, which is consistent with our observations (Fig. 2; see [131] for recent results demonstrating a similar phenomenon in grasses infected by Epichloë endophytes).

To account for any adverse effects of seed coat removal, we planted control seeds alongside sterile agar or agar inoculated with *A. fulva*. This technique was successful as shown by culturing results (Fig. S2), swainsonine concentrations (Fig. S4), and sequencing output (Fig. S8). The results we observed from control plants mirrored those from treated plants, except for foliar C and N concentrations, which were more variable among controls. Most manipulative studies of vertically-transmitted fungal endophytes reduce fungi within seeds through either heat treatment (e.g. [132]), long-term storage (e.g. [12]), or fungicide application (e.g. [133, 134]). While studies manipulating endophytes via these techniques have been of critical importance, it is possible that these treatments could have undesirable consequences that obscure effects of endophyte reduction. Consequently, we suggest others consider the approach we use here when seeds from endophyte-free plants are not available.

### Conclusion

Our results highlight the simultaneous, contrasting effects foliar endophytes can have on their hosts and the complexity of microbe-microbe interactions. For instance, pathogen pressure was reduced by *A. fulva* presence, but plants colonized by this fungus were smaller. Moreover, inoculation of hosts with rare endophytes led to smaller host size and reduced foliar %N, but only when *A. fulva* was not present. Thus, *A. fulva* moderates the effects of both rare endophytes and a dominant pathogen on the host. Our study affirms the recent emphasis on dominant, prevalent microbes as a tractable way to parse the complexity of the foliar microbiome. However, we suggest that rare endophytic taxa, occupying only small portions of their hosts, should not be neglected as their ecological consequences could exceed expectations.

## Acknowledgments and conflict of interest statement

We declare no conflicts of interest. Funding was provided by a National Science Foundation (NSF) Doctoral Dissertation Grant awarded to JGH. JGH and CAB were supported by the Microbial Ecology Collaborative with funding from NSF award #EPS-1655726. Computing was performed in the Teton Computing Environment at the Advanced Research Computing Center, University of Wyoming, Laramie (https://doi.org/10.15786/M2FY47).

## Data availability

All scripts, plant trait data, and processed sequence data are available at: https://github.com/JHarrisonEcoEvo/ASLE_microbiome_manipulation.git Raw data are hosted by the University of Wyoming at: https://doi.org/10.15786/r9xy-6x03

## Supplementary Material

### Library preparation details

A two-step PCR was used to prepare sequencing libraries. In the first step, the locus of interest was amplified and in a subsequent step Illumina adapters were added to amplicons. Both 16s and ITS libraries were created using triplicate 20 μl PCR reactions (1 μl each of forward and reverse primers, 10 μl of NEBNext 2X Master Mix [New England BioLabs, Ipswich, MA], 7μl of water, and 1 μl of template [~9 ng of DNA]). PCR conditions for both markers were identical. Conditions for the first round of amplification were as follows: an initial denaturation at 98°C for 30 s, followed by 12 cycles of 98°C for 30 s, 62°C for 30 s, and 72°C for 30 s, and a final 5 min extension at 72°C. Amplicons were cleaned using AMPure beads (0.8x; Beckman Coulter, Indianapolis, IN, U.S.A) prior to the second round of PCR. Reaction volume for the second round of PCR was 30 μl (15 μl NEBNext 2X Master Mix, 5 μl primers, 10 μl of product from the first round of PCR). Conditions for this round were the same as the first round, except only 7 cycles were conducted. Amplicons were cleaned again using AMPure beads and qPCR used to determine concentration. Amplicons were pooled in equimolar fashion, quality determined via a Bio-Analyzer (Agilent, Santa Clara, CA, U.S.A), and sequenced.

### Sequencing of mock community

For ITS data obtained from sequencing the mock community, 1 ESV corresponded with *Cryptococcus neoformans*, and 3 ESVs with *Saccharomyces cerevisiae*, the two fungi expected to occur in the ZymoBIOMICS mock community (Zymo Research, Irvine, CA. U.S.A.). An additional fungal ESV was present but only had two reads, and so is likely a very rare lab contaminant or sequencing/PCR artifact. For 97% OTUs, one OTU matched *C. neoformans* and one OTU matched *S. cerevisiae.* We observed 13 ESVs in the 16s data from the mock community that were present at > 14 reads. This is slightly higher than the eight taxa expected. An additional two ESVs were present with < 14 reads. For 97% OTUS, we recovered eight OTUs as expected. Bacterial taxa in the mock community included: *Listeria monocytogenes, Pseudomonas aeruginosa, Bacillus subtilis, Escherichia coli, Salmonella enterica, Lactobacillus fermentum, Enterococcus faecalis* and *Staphylococcus aureus*. Our taxonomic hypothesis generation method accurately captured the genus of each of these taxa for both ESVs and 97% OTUs, and in most cases species level resolution was possible.

Despite the slight overestimation of mock community richness when using ESVs, we choose to use ESVs for our analyses because of the logistical and biological benefits they provide compared to 97% OTUs. First, much ecologically and evolutionarily important genetic information exits below the species level and ESVs are more likely to reveal such variation. Indeed, such cryptic variation could explain the overestimation of richness in the ZymoBIOMICS mock community that we observed. Additionally, Harrison et al.[56] found certain ESVs of *A. fulva*, the heritable fungus within *A. lentiginosus*, were more associated with swainsonine variation than other ESVs. Second, many technical concerns arise from the use of 97% OTUs. For example, the lengths of sequences determine how many bases need to differ for those sequences to diverge at 3% or greater. In other words, the denominator in percentage calculations depends on sequence length, which likely differs across phylogeny, thus 97% do not correspond to the same amount of genetic variation for each taxon. Other logistical concerns exist and, for further discussion, the reader is directed to [81]. *Sequencing results*

A total of 2,207,373 reads were generated through sequencing of the ITS1 library. Of these 2,064,256 merged and 2,020,983 mapped to ESVs used to generate an ESV table. The majority of these reads were from the *A. lentiginosus* host, which was to be expected given that plants were not grown outdoors for more than several months. A total of 115,860 reads from 11 fungal ESVs were obtained from focal plants.

Sequencing of the 16s library generated 2,515,615 reads, of which 2,364,256 merged. Of these 2,136,863 mapped to ESVs used to generate an ESV table. The majority of these reads were host chloroplast DNA. In total, 1,496 reads from 53 bacterial ESVs were recovered from focal plant tissues. This low number of reads is not due to issues with library preparation and sequencing as 31,376 bacterial reads were obtained from inoculum samples, but was instead due to the very high relative abundance of chloroplast DNA to endophytic bacteria.

**Table S1:**
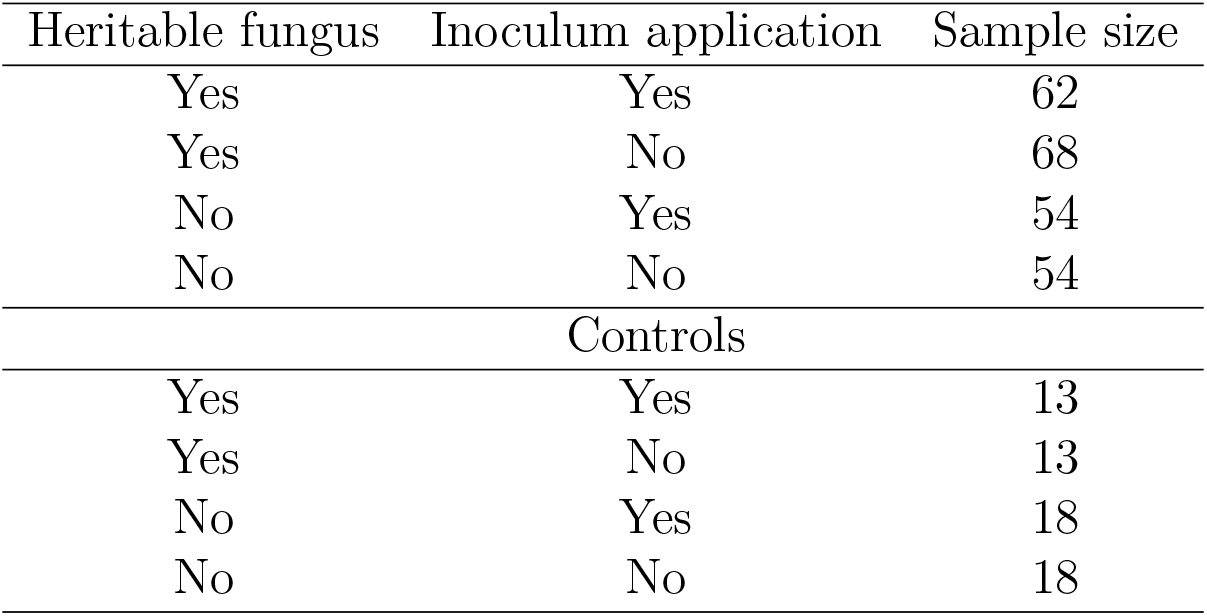
Number of *A. lentiginosus* plants installed in field experiment for each treatment group. Treatments were applied to plants in a full factorial design. Treatments included reduction of the relative abundance of the vertically-transmitted fungal endophyte (through embryo excision) and foliar application of locally-sourced, endophyte inocula. To control for effects of seed coat removal, embryos were excised from seed coats and planted alongside agar that was either sterile or contained *A. fulva*.

**Table S2:** Trait values for *A. lentiginosus* individuals in each treatment group. Values shown are means of posterior probability distributions of the mean for each trait with 95% credible intervals in parentheses.

|  | Treated – | Treated + | Untreated – | Untreated + |
| --- | --- | --- | --- | --- |
| Size (cm<sup>3</sup>) | 10028.92 (10000.52,10058.05) | 14631.37 (14601.26,14662.02) | 23880.19 (23849.97,23910.29) | 7423.03 (7394.08,7452.41) |
| Leaves | 28.69 (21.21,35.81) | 27.3 (17.35,37.41) | 50.46 (33.38,67.69) | 25.81 (11.59,39.66) |
| Leaflet area (cm<sup>2</sup>) | 0.49 (0.46,0.52) | 0.53 (0.49,0.57) | 0.53 (0.49,0.56) | 0.51 (0.48,0.54) |
| SLA (cm<sup>2</sup>mg<sup>-1</sup>) | 0.12 (0.11,0.13) | 0.12 (0.11,0.13) | 0.12 (0.11,0.13) | 0.12 (0.11,0.13) |
| delta N15 | 0.52 (-0.02,1.03) | 1.33 (0.79,1.88) | 0.84 (0.09,1.58) | 1.15 (-0.15,2.33) |
| % swainsonine | 0 (0,0) | 0.03 (0.01,0.04) | 0 (0,0) | 0.04 (0.02,0.06) |
| % N | 0.87 (0.79,0.95) | 0.93 (0.81,1.06) | 1.03 (0.92,1.13) | 0.88 (0.74,1.02) |
| % C | 12.32 (11.87,12.76) | 12.46 (11.86,13.06) | 11.92 (11.3,12.52) | 12.26 (11.62,12.88) |
| NDFA | 62.14 (55.19,69.15) | 51.11 (43.55,59.04) | 57.56 (46.7,67.5) | 53.43 (36.49,69.31) |

**Table S3:** Effect of treatment on microbial relative abundance. Values shown are the number of microbial taxa differing in relative abundance for each pairwise comparison. The upper half of the table (shaded) describes results for fungi (9 taxa considered), and the bottom half of the table describes results for bacteria (53 taxa considered). ESVs corresponding to *A. fulva* genotypes were omitted during analysis, however these ESVs were also influenced by treatment (see Fig. S1). Differences in relative abundance were determined through a hierarchical Bayesian modelling approach (see main text).

|  | Asle +<br>Inoculum + | Asle +<br>Inoculum - | Asle -<br>Inoculum + | Asle -<br>Inoculum - |
| --- | --- | --- | --- | --- |
| Asle +<br>Inoculum + | - | 1 | 0 | 1 |
| Asle +<br>Inoculum - | 0 | - | 4 | 4 |
| Asle -<br>Inoculum + | 0 | 0 | - | 0 |
| Asle -<br>Inoculum - | 0 | 0 | 0 | - |

**Table S4:** Associations between plant traits and species equivalencies of Shannon’s entropy for fungi (shaded, top rows) and bacteria (unshaded, bottom rows). Values in cells are the mean of posterior probability distributions (PPDs) of beta coefficient estimates from a hierarchical linear model calculated in a Bayesian framework. Values in parentheses denote the proportion of each PPD greater than zero, which is the certainty of a non-zero effect of the covariate on the response.

| Treatment | Intercept | Plant<br>volume | SLA | %N | %C | NDFA | Swainsonine |
| --- | --- | --- | --- | --- | --- | --- | --- |
| Asle –<br>Inoculum + | **1.16**<br>(100%) | -0.02<br>(48.1%) | 0.02<br>(56.2%) | 0.01<br>(54.0%) | -0.01<br>(45.9%) | 0.00<br>(48.1%) | – |
| Asle +<br>Inoculum + | **1.15**<br>(100%) | **-0.02**<br>(3.5%) | 0.00<br>(25.9%) | 0.00<br>(64.3%) | 0.00<br>(30.1%) | 0.01<br>(88.8%) | 0.12<br>(57.7%) |
| Asle –<br>Inoculum – | **1.17**<br>(100%) | 0.00<br>(45.5%) | -0.01<br>(15.8%) | 0.01<br>(77.2%) | 0.00<br>(64.1%) | 0.00<br>(32.5%) | – |
| Asle +<br>Inoculum – | **1.14**<br>(100%) | 0.01<br>(89.8%) | 0.00<br>(41.7%) | -0.02<br>(6.9%) | 0.00<br>(57.4%) | 0.00<br>(48.5%) | 0.03<br>(70.2%) |
| Asle –<br>Inoculum + | 1.15<br>(100%) | **-0.02**<br>(2.8%) | 0.00<br>(26.1%) | 0.00<br>(66.2%) | 0.00<br>(28.6%) | -0.01<br>(49.4%) | – |
| Asle +<br>Inoculum + | **1.17**<br>(100%) | 0.00<br>(45.4%) | -0.01<br>(18.6%) | 0.01<br>(77.6%) | 0.00<br>(64.5%) | 0.01<br>(69.2%) | 0.12<br>(54.9%) |
| Asle –<br>Inoculum – | **1.14**<br>(100%) | 0.01<br>(85.7%) | 0.00<br>(39.9%) | -0.02<br>(11.6%) | 0.00<br>(54.5%) | -0.00<br>(35.5%) | – |
| Asle +<br>Inoculum – | 1.2 (100%) | 0.03<br>(65.3%) | 0.00<br>(54.8%) | -0.01<br>(40.2%) | **-0.07**<br>(4.5%) | 0.00<br>(50.2%) | 0.01<br>(51.5%) |

**Table S5:** Associations between plant traits and species equivalencies of Simpson’s diversity for fungi (shaded, top rows) and bacteria (unshaded, bottom rows). Values in cells are the mean of posterior probability distributions (PPDs) of beta coefficient estimates from a hierarchical linear model calculated in a Bayesian framework. Values in parentheses denote the proportion of each PPD greater than zero, which is the certainty of a non-zero effect of the covariate on the response.

| Treatment | Intercept | Plant<br>volume | SLA | %N | %C | NDFA | Swainsonine |
| --- | --- | --- | --- | --- | --- | --- | --- |
| Asle –<br>Inoculum + | **23.83**<br>(100%) | -2.07<br>(36.9%) | 0.77<br>(84.8%) | -0.22<br>(37.6%) | 0.6<br>(73.5%) | -0.48<br>(32.6%) | – |
| Asle +<br>Inoculum + | **25.13**<br>(100%) | **4.11**<br>(99.5%) | 0.83<br>(93.0%) | -0.43<br>(26.2%) | 0.45<br>(75.3%) | -0.62<br>(18.1%) | -1.57<br>(40.3%) |
| Asle –<br>Inoculum – | **21.21**<br>(100%) | -0.13<br>(38.3%) | **1.03**<br>(98.3%) | -0.57<br>(13.2%) | -0.11<br>(41.1%) | 0.10<br>(59.2%) | – |
| Asle +<br>Inoculum – | **25.88**<br>(100%) | -0.04<br>(48.8%) | **1.11**<br>(92.7%) | 0.16<br>(50.0%) | 0.58<br>(72.8%) | -0.24<br>(39.0%) | **-1.29**<br>(0.0%) |
| Asle –<br>Inoculum + | **25.1**<br>(100%) | **4.14**<br>(99.8%) | 0.86<br>(94.3%) | -0.52<br>(21.2%) | 0.51<br>(78.8%) | -0.66<br>(15.2%) | – |
| Asle +<br>Inoculum + | 21.26<br>(100%) | -0.12<br>(38.2%) | 1.10<br>(98.6%) | -0.61<br>(9.7%) | -0.07<br>(44.6%) | 0.05<br>(54.4%) | 0.13<br>(50.8%) |
| Asle –<br>Inoculum – | 25.94<br>(100%) | 0.11<br>(53.3%) | 1.15<br>(93.0%) | 0.12<br>(47.6%) | 0.77<br>(74.6%) | -0.28<br>(35.8%) | – |
| Asle +<br>Inoculum – | 22.11<br>(100%) | 0.04<br>(51.2%) | 0.99<br>(89.9%) | -0.14<br>(39.8%) | 93 (78.2%) | -0.13<br>(42.1%) | 2.7<br>(84.4%) |

**Figure S1:**
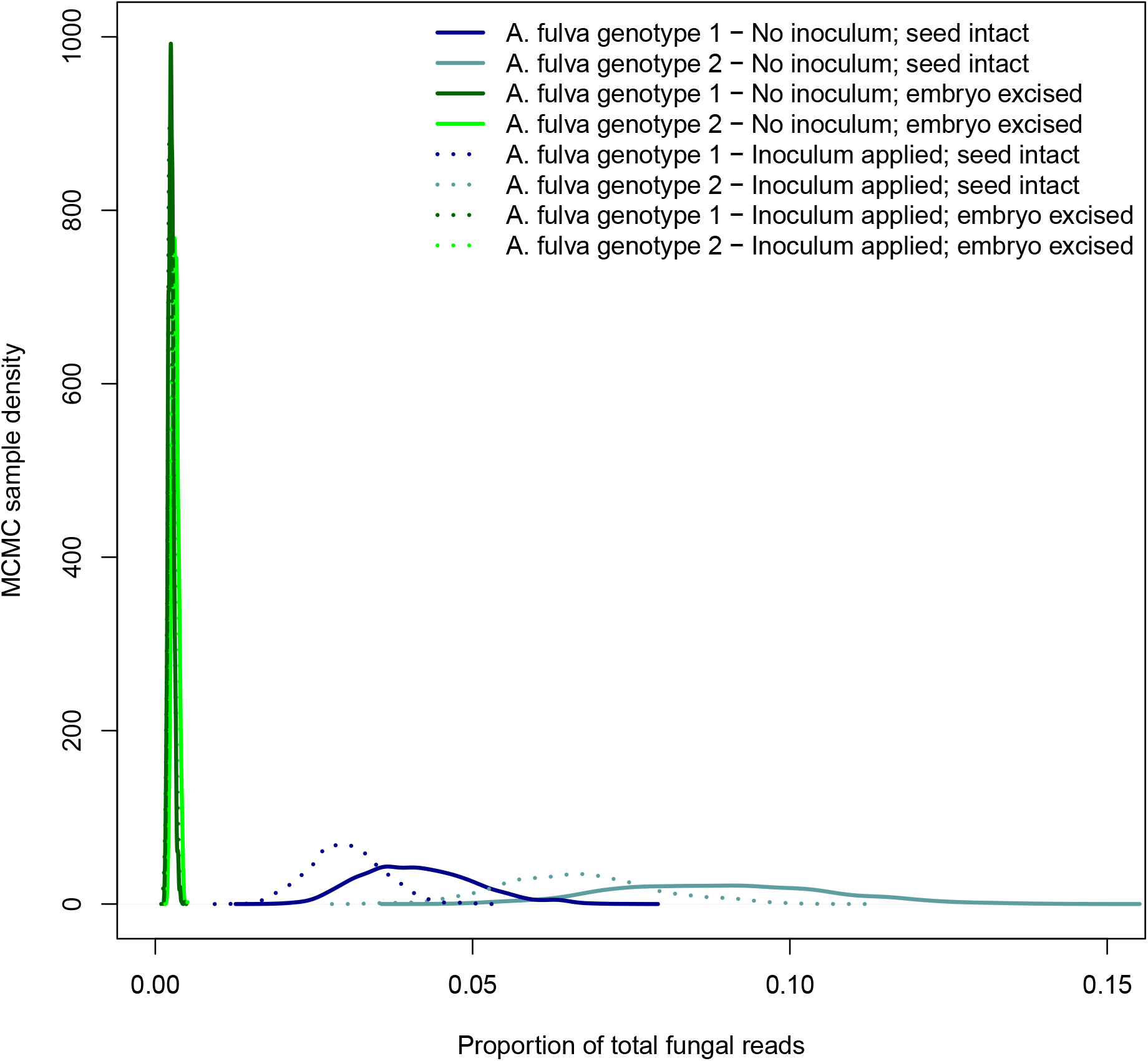
Estimated proportion of fungal reads that were assigned to *A. fulva*. Estimates shown are posterior probability distributions from the hierarchical Bayesian analysis (see main text). Two genotypes of *A. fulva* were identified (two ESVs). Culturing revealed these two genotypes exhibited different phenotypes. Plants reared from embryos excised from the seed coat had lower relative abundance of *A. fulva*, as shown through sequencing and culturing (also see Fig. S2).

**Figure S2:**
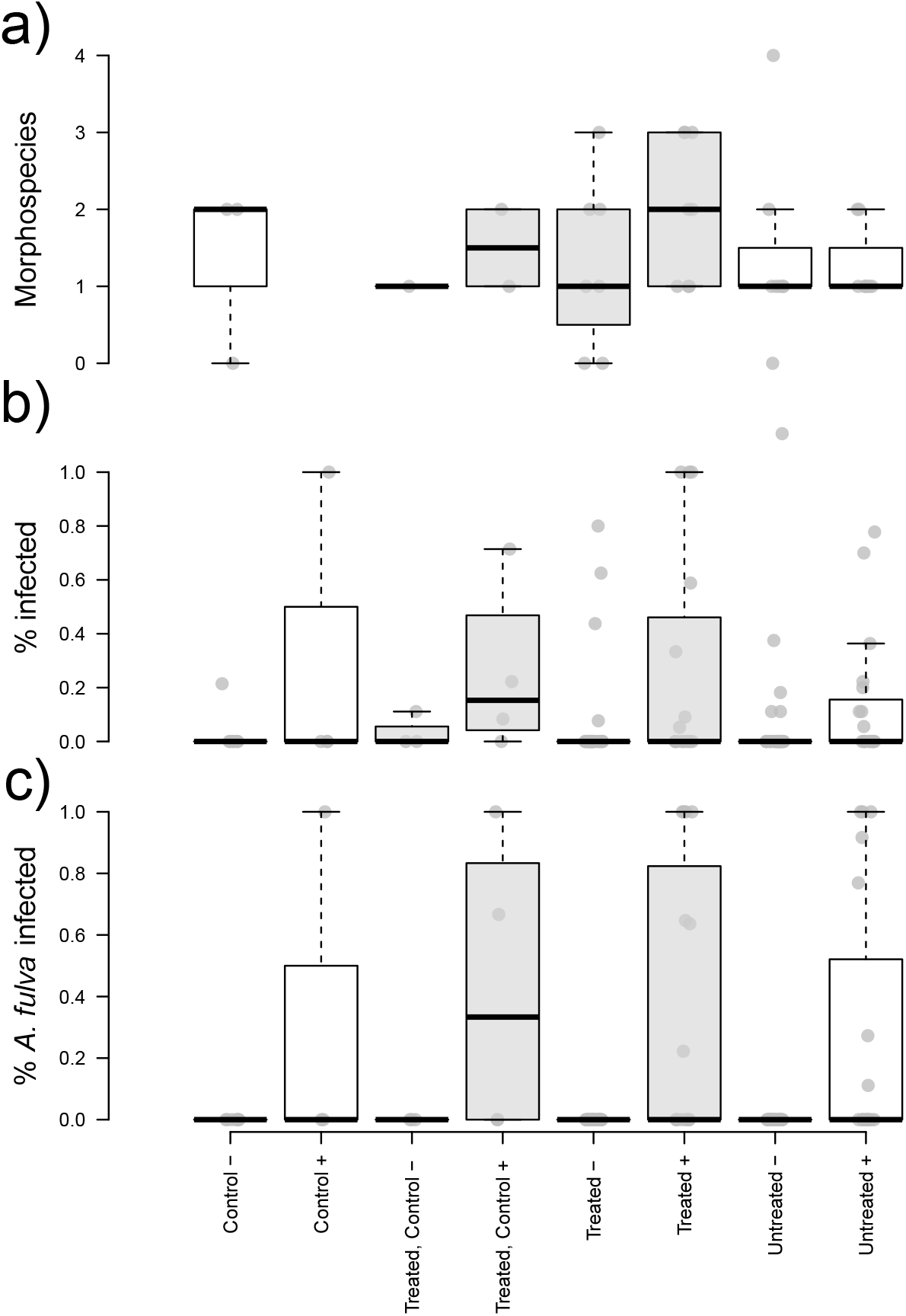
Results from culturing of *A. lentiginosus* foliar tissue. + and - symbols denote treatment to reduce the relative abundance of the vertically-transmitted fungus, *A. fulva.* Treatment groups that received inoculum mixture are prefixed with “Treated” and boxes shaded. Control treatment involved removal of the seed coat and planting alongside sterile agar or agar containing *A. fulva*. Panel a shows the number of morphospecies observed for each treatment group, not counting *A. fulva*. Panel b shows the percentage of foliar tissue segments colonized by non-*A. fulva* microbes. Panel c shows the percentage of tissue segments colonized by *A. fulva*. Correct identification of *A. fulva* cultures was confirmed via sequencing. Boxplots summarize the data and describe interquartile range with a horizontal line denoting the median. Whiskers extend to the 10th and 90th percentiles.

**Figure S3:**
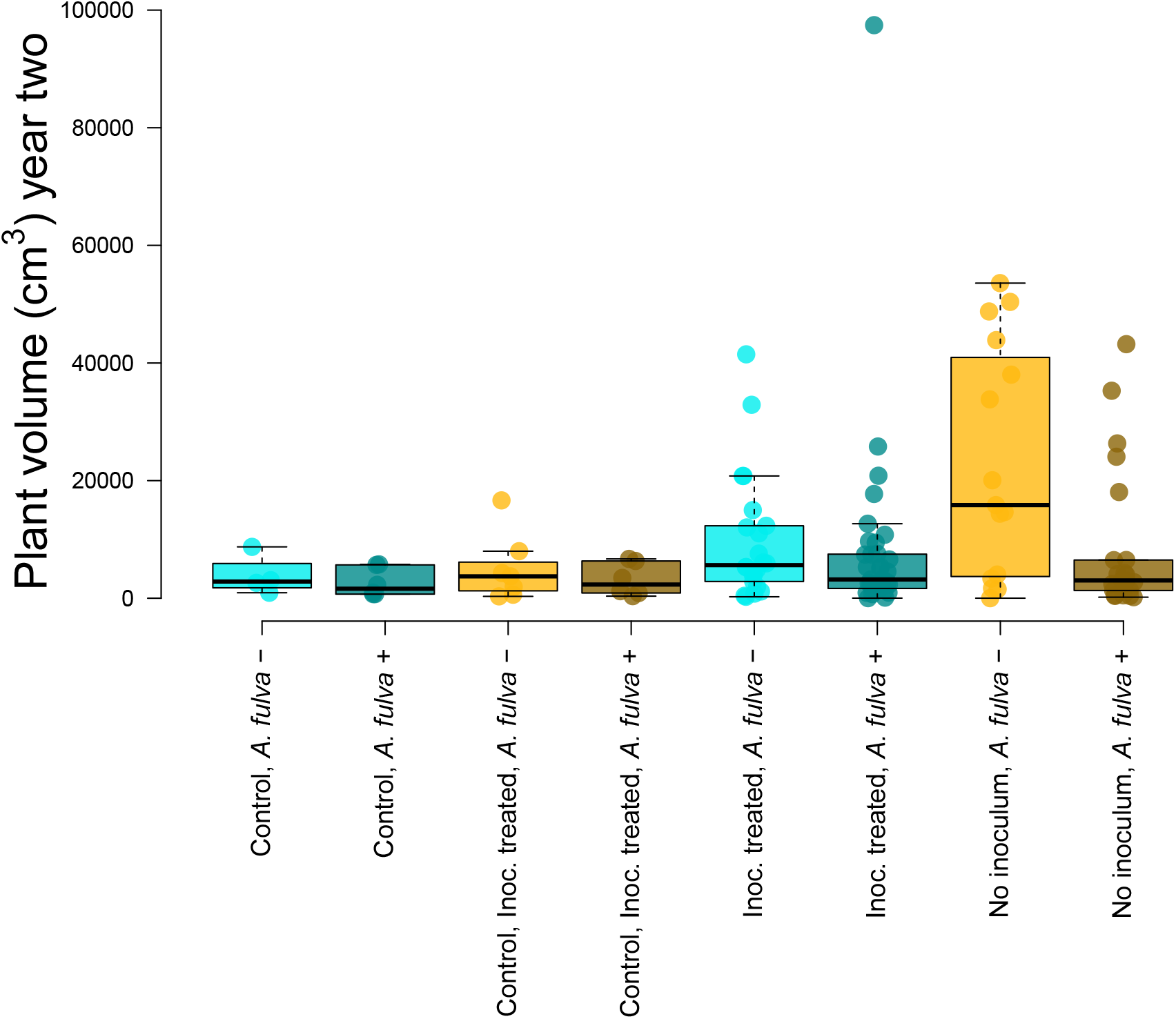
Size of *A. lentiginosus* plants in the year following initial experiment set up and sample collection. The smaller size of *A. fulva* colonized plants is apparent and mirrors what was observed during the year the experiment was conducted. Boxplots summarize the data and describe interquartile range with a horizontal line denoting the median. Whiskers extend to the 10th and 90th percentiles.

**Figure S4:**
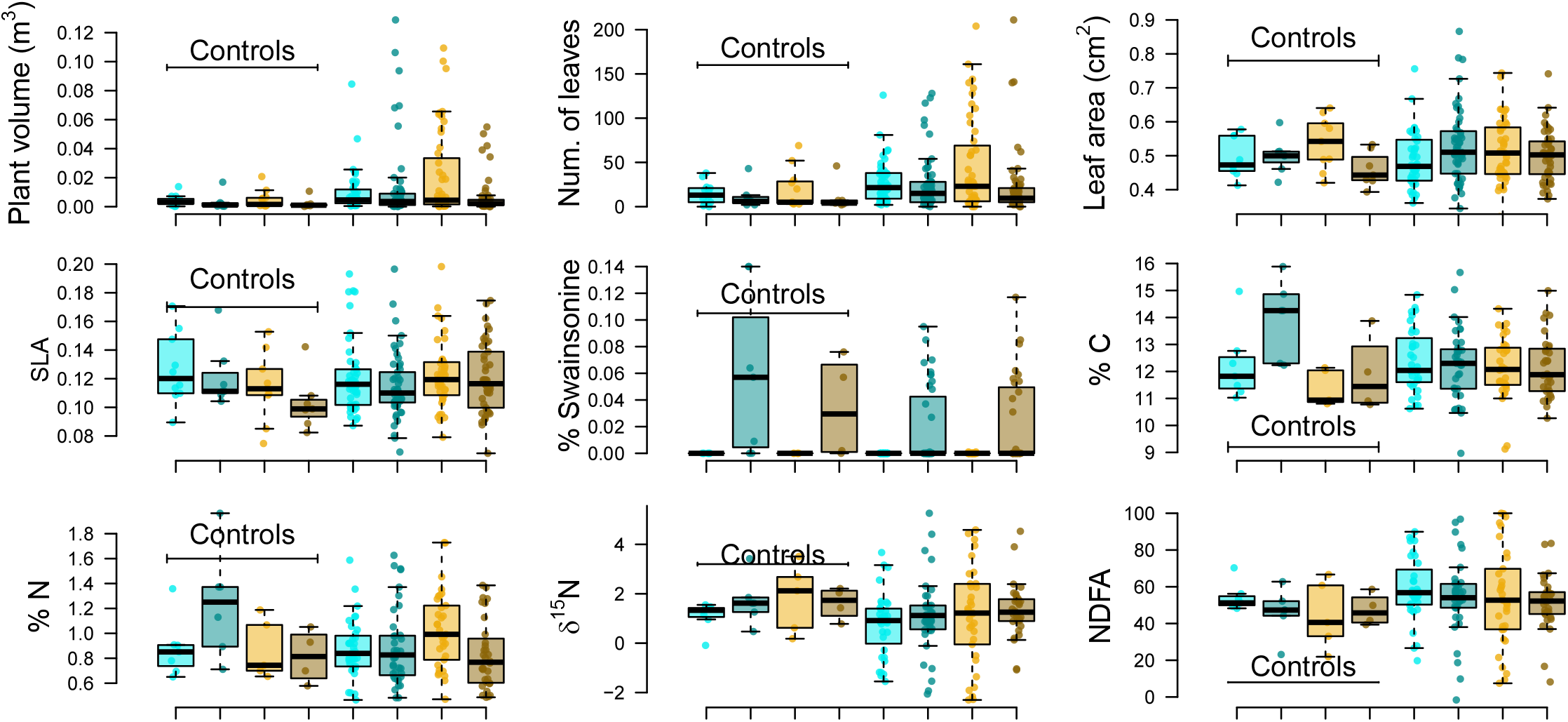
Influence of treatment on plant traits. This figures mirrors Fig. 1, except that control treatment groups are also shown. % C and N is the % of dry mass of foliar tissue composed of either element. For details of treatments and trait measurements see the main text. Boxplots summarize the data and describe interquartile range with a horizontal line denoting the median. Whiskers extend to the 10th and 90th percentiles.

**Figure S5:**
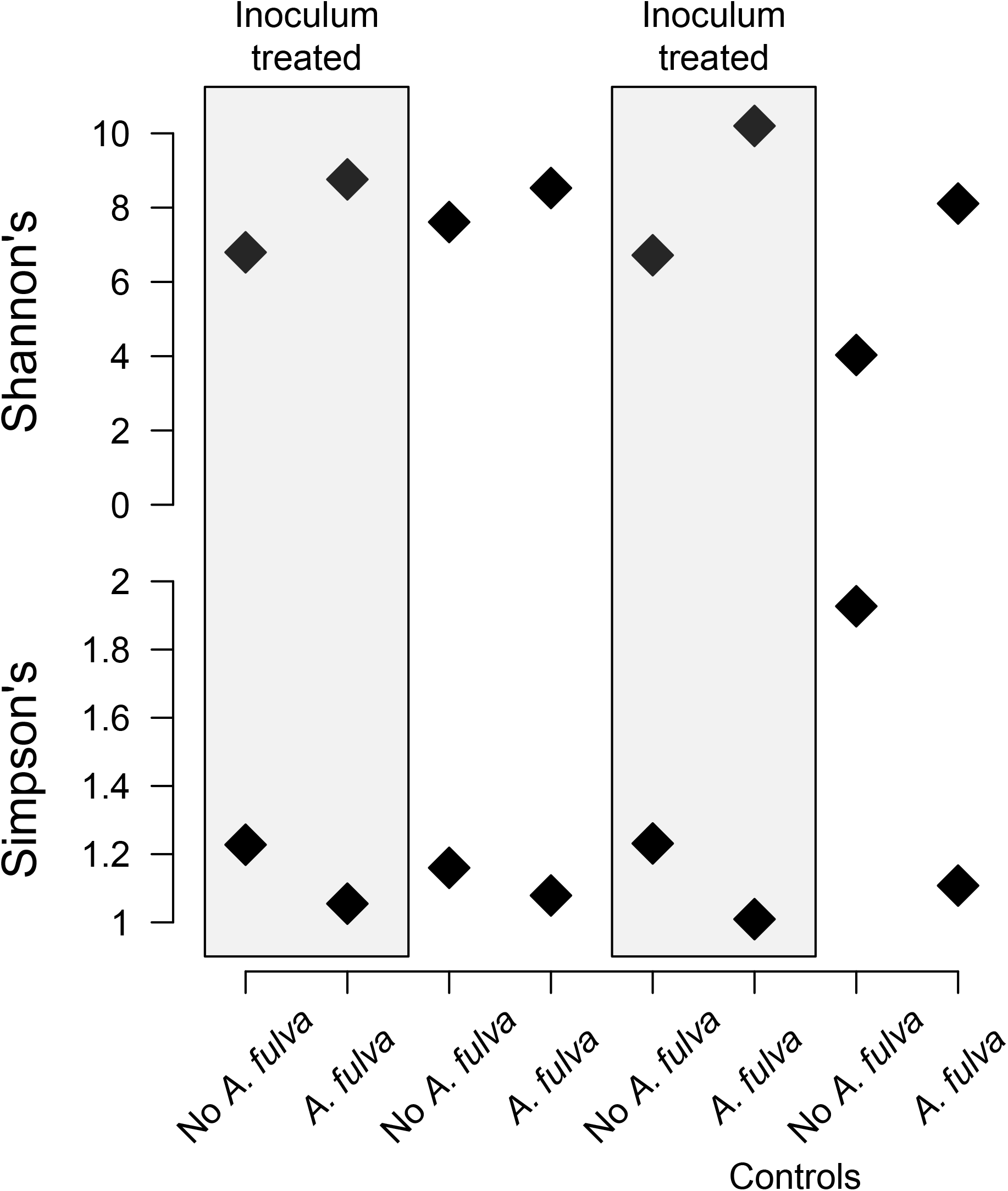
Influence of treatment on fungal and bacterial diversity. This figure shows the same data as Fig. 2, but also includes control treatment groups.

**Figure S6:**
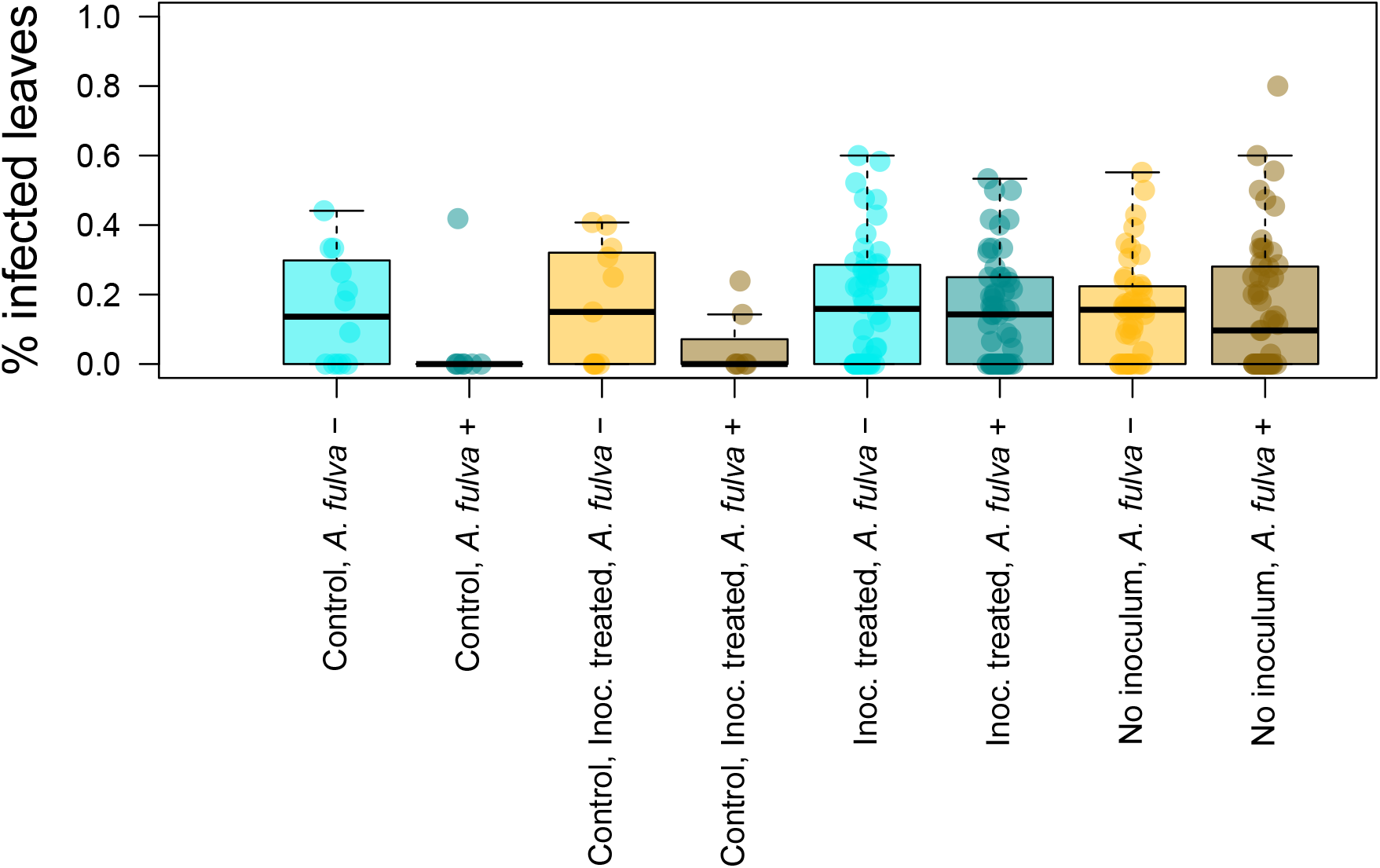
Differences in powdery mildew infection among treatment groups. The percentage of infected leaves observed is shown on the y- axis. Infection was determined visually (a whitish growth on leaf surfaces). Boxplots summarize the data and describe interquartile range with a horizontal line denoting the median. Whiskers extend to the 10th and 90th percentiles. Posterior probability distributions (PPDs) of the mean difference between groups differed for control seedlings with high certainty (>80% of the PPDs did not overlap). Much more overlap was observed for non-control plants, though the directionality of effect was consistent for all treatment groups. Specifically, *A. fulva* colonization was associated with reduced mildew presence.

**Figure S7:**
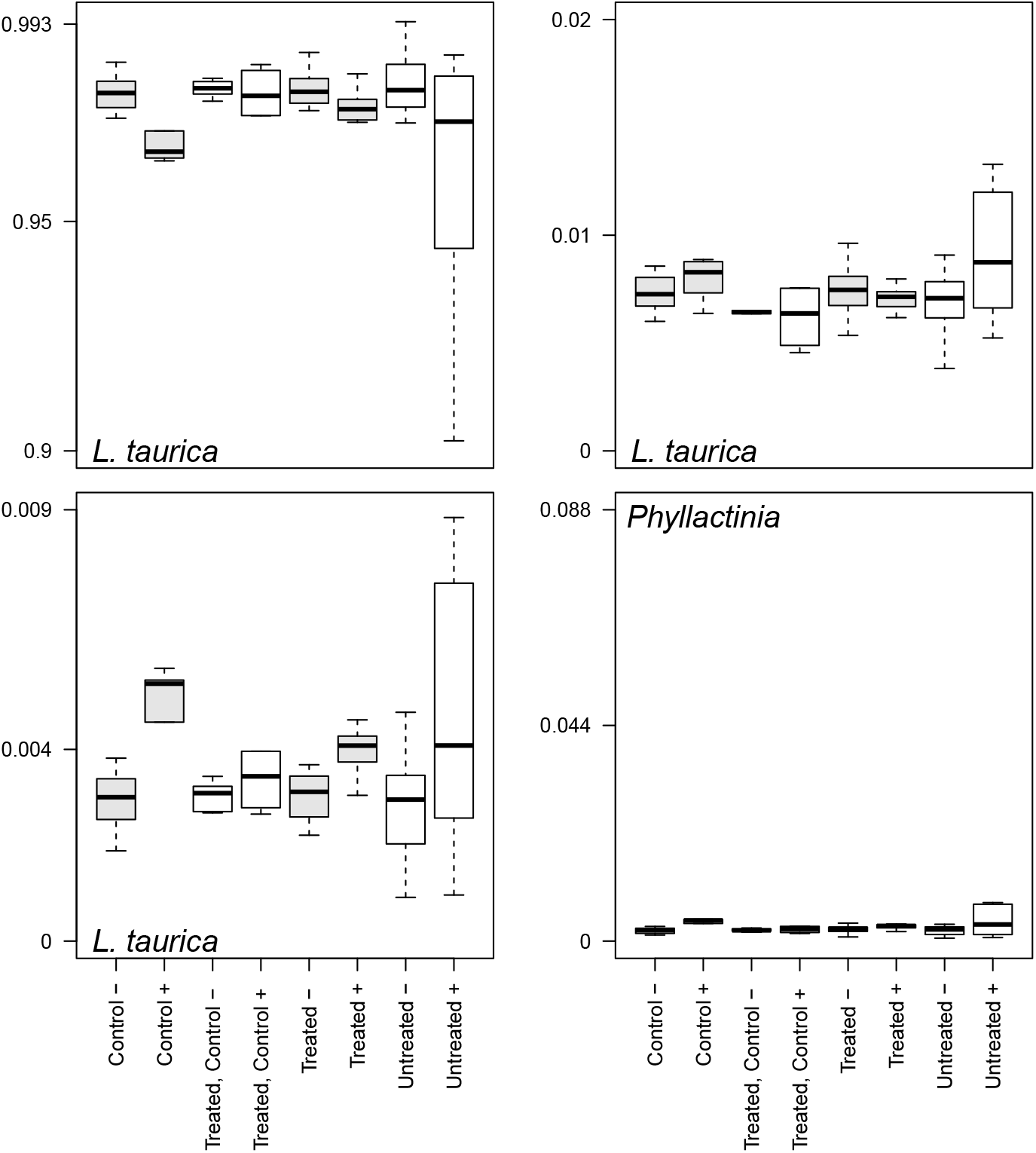
Differences among treatment groups in the relative abundance of abundant fungal endophyte taxa. The y axis is proportion of reads from the sample composed of the taxon shown (means of posterior probability distributions of *p* parameters from multinomial distribution). Boxplots show the interquartile range with a horizontal line denoting the median. WWhiskers extend to the 10th and 90th percentiles. There were several genotypes of *L. taurica* represented in the sequence data, though the dominant genotype (upper left panel) was much more abundant than the rarer forms. In most cases, the presence of *A. fulva* influenced the relative abundance of these fungal taxa (statistical tests via the hierarchical Bayesian approach described in the main text)

**Figure S8:**
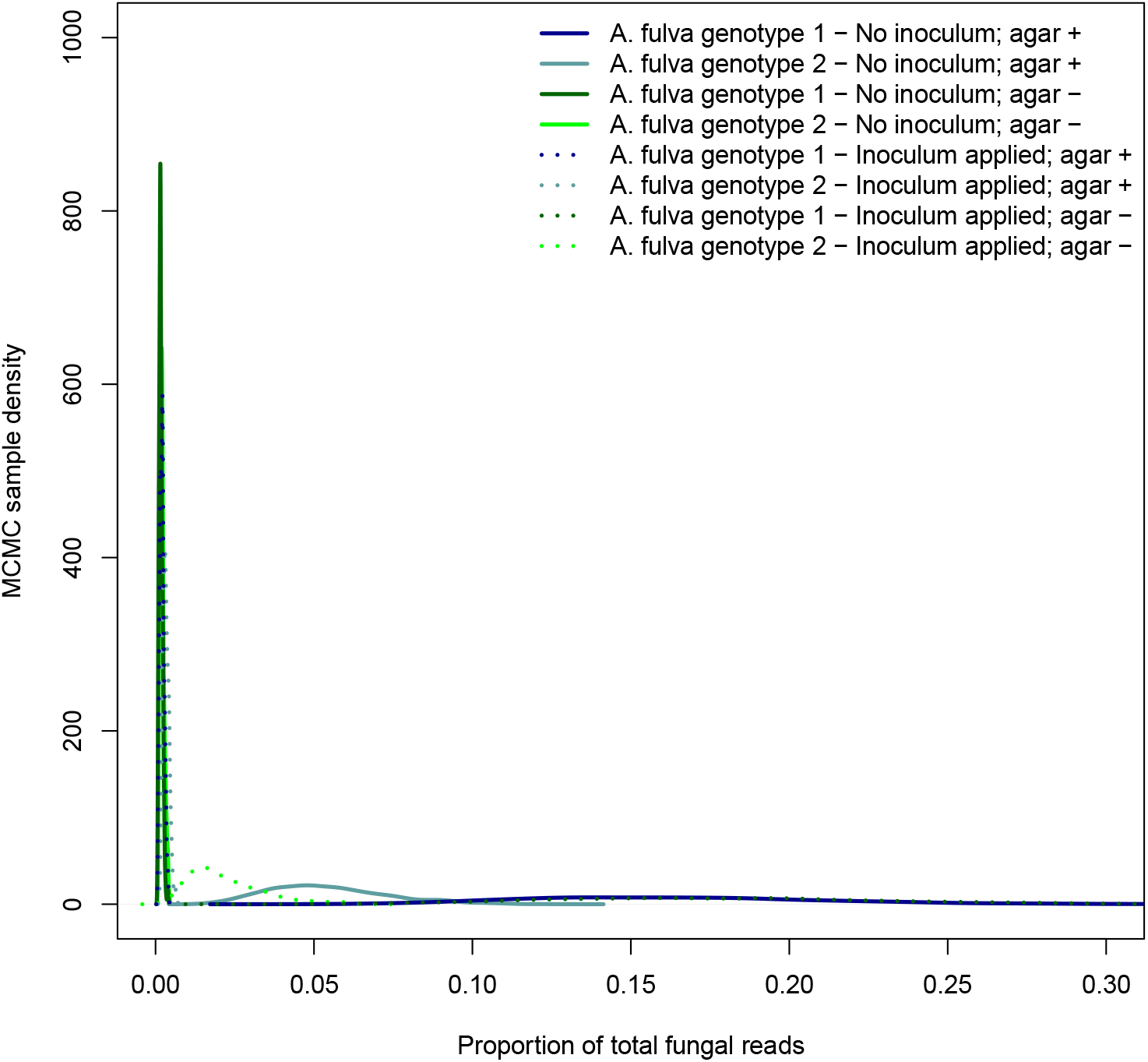
Estimated proportion of fungal reads that were assigned to *A. fulva* and recovered from control plants. These plants were grown from seeds planted alongside a chunk of sterile agar or *A. fulva* colonized agar. Plants reared alongside sterile agar had lower proportions of *A. fulva* than plants reared alongside *A. fulva* colonized agar. This result was also confirmed via swainsonine analysis and culturing (Fig. S2, S4). Estimates shown are posterior probability distributions from the hierarchical Bayesian analysis (see main text). Two genotypes of *A. fulva* were identified (two ESVs). Culturing revealed these two genotypes exhibited different phenotypes.

## References

1. Ryan, R. P., Germaine, K., Franks, A., Ryan, D. J. & Dowling, D. N. Bacterial endophytes: recent developments and applications. en. FEMS Microbiology Letters 278, 1–9. ISSN: 0378-1097, 1574-6968 (Jan. 2008).

2. Rodriguez, R., White Jr, J., Arnold, A. & Redman, R. Fungal endophytes: diversity and functional roles. New Phytologist 182, 314–330. ISSN: 1469-8137 (2009).

3. Griffin, E. A. & Carson, W. P. The ecology and natural history of foliar bacteria with a focus on tropical forests and agroecosystems. en. The Botanical Review 81, 105–149. ISSN: 1874-9372 (June 2015).

4. Carroll, G. Fungal endophytes in stems and leaves: from latent pathogen to mutualistic symbiont. en. Ecology 69, 2–9. ISSN: 1939-9170 (1988).

5. Wilson, D. Endophyte: the evolution of a term, and clarification of its use and definition. Oikos, 274–276. ISSN: 0030-1299 (1995).

6. Porras-Alfaro, A. & Bayman, P. Hidden fungi, emergent properties: endophytes and microbiomes. Phytopathology 49, 291 (2011).

7. Hardoim, P. R. et al. The hidden world within plants: ecological and evolutionary considerations for defining functioning of microbial endophytes. en. Microbiology and Molecular Biology Reviews 79, 293–320. ISSN: 1092-2172, 1098-5557 (Sept. 2015).

8. Petrini, O. Fungal endophytes of tree leaves en. in Microbial Ecology of Leaves (eds Andrews, J. H. & Hirano, S. S.) (Springer New York, 1991), 179–197. ISBN: 978-14612-3168-4.

9. Arnold, A. E. & Lutzoni, F. Diversity and host range of foliar fungal endophytes: are tropical leaves biodiversity hotspots? en. Ecology 88, 541–549. ISSN: 1939-9170 (Mar. 2007).

10. Clay, K. & Schardl, C. Evolutionary origins and ecological consequences of endophyte symbiosis with grasses. The American Naturalist 160, S99–S127. ISSN: 0003-0147 (2002).

11. Rudgers, J. A. et al. A fungus among us: broad patterns of endophyte distribution in the grasses. en. Ecology 90, 1531–1539. ISSN: 1939-9170 (June 2009).

12. Clay, K. & Holah, J. Fungal endophyte symbiosis and plant diversity in successional fields. en. Science 285, 1742–1744. ISSN: 0036-8075, 1095-9203 (Sept. 1999).

13. Afkhami, M. E. & Strauss, S. Y. Native fungal endophytes suppress an exotic dominant and increase plant diversity over small and large spatial scales. en. Ecology 97, 1159–1169. ISSN: 1939-9170 (2016).

14. Rudgers, J. A. & Clay, K. Endophyte symbiosis with tall fescue: how strong are the impacts on communities and ecosystems? Fungal Biology Reviews 21, 107–124 (2007).

15. Rudgers, J. A. & Clay, K. An invasive plant–fungal mutualism reduces arthropod diversity. en. Ecology Letters 11, 831–840. ISSN: 1461-0248 (2008).

16. Gorischek, A. M., Afkhami, M. E., Seifert, E. K. & Rudgers, J. A. Fungal symbionts as manipulators of plant reproductive biology. The American Naturalist 181, 562–570. ISSN: 0003-0147 (Apr. 2013).

17. Malloch, D. & Blackwell, M. in The fungal community: its organization and role in the ecosystem Second Edition, 147–171 (Marcel Dekker, Inc, New York, NY, 1992).

18. Devarajan, P. & Suryanarayanan, T. Evidence for the role of phytophagous insects in dispersal of non-grass fungal endophytes. Fungal Diversity 23, 111–119 (Oct. 2006).

19. Hodgson, S. et al. Vertical transmission of fungal endophytes is widespread in forbs. en. Ecology and Evolution 4, 1199–1208. ISSN: 2045-7758 (Apr. 2014).

20. Newcombe, G., Harding, A., Ridout, M. & Busby, P. A hypothetical bottleneck in the plant microbiome. English. Frontiers in Microbiology 9. ISSN: 1664-302X. doi:10.3389/fmicb.2018.01645. https://www.frontiersin.org/articles/10.3389/fmicb.2018.01645/abstract (2018) (2018).

21. Lodge, D. J., Fisher, P. & Sutton, B. Endophytic fungi of *Manilkara bidentata* leaves in Puerto Rico. Mycologia, 733–738. ISSN: 0027-5514 (1996).

22. Paine, R. T. A note on trophic complexity and community stability. The American Naturalist 103, 91–93. ISSN: 0003-0147 (Jan. 1969).

23. Davic, R. D. Linking keystone species and functional groups: a new operational definition of the keystone species concept. Conservation Ecology 7. ISSN: 1195-5449. https://www.jstor.org/stable/26271938 (2019) (2003).

24. Jenkins, S. H. & Busher, P. E. *Castor canadensis*. en. Mammalian Species, 1–8. ISSN: 0076-3519 (June 1979).

25. Rosell, F., Bozsér, O., Collen, P. & Parker, H. Ecological impact of beavers *Castor fiber* and *Castor canadensis* and their ability to modify ecosystems. en. Mammal Review 35, 248–276. ISSN: 1365-2907 (2005).

26. Hajishengallis, G., Darveau, R. P. & Curtis, M. A. The keystone-pathogen hypothesis. en. Nature Reviews Microbiology 10, 717–725. ISSN: 1740-1526, 1740-1534 (Oct. 2012).

27. Jousset, A. et al. Where less may be more: how the rare biosphere pulls ecosystems strings. en. The ISME Journal 11, 853–862. ISSN: 1751-7370 (Apr. 2017).

28. Hassani, M. A., Durán, P. & Hacquard, S. Microbial interactions within the plant holobiont. Microbiome 6, 58. ISSN: 2049-2618 (Mar. 2018).

29. Rockman, M. V. The QTN program and the alleles that matter for evolution: all that’s gold does not glitter. en. Evolution 66, 1–17. ISSN: 1558-5646 (2012).

30. Kloepper, J. W. & Ryu, C.-M. en. in Microbial Root Endophytes (eds Schulz, B. J. E., Boyle, C. J. C. & Sieber, T. N.) 33–52 (Springer Berlin Heidelberg, Berlin, Heidelberg, 2006). ISBN: 978-3-540-33526-9. doi:10.1007/3-540-33526-9_3. https://doi.org/10.1007/3-540-33526-9_3 (2019).

31. Beckers, G. J. & Conrath, U. Priming for stress resistance: from the lab to the field. Current Opinion in Plant Biology. Special Issue on Biotic Interactions 10, 425–431. ISSN: 1369-5266 (Aug. 2007).

32. Conn, V. M., Walker, A. R. & Franco, C. M. M. Endophytic Actinobacteria induce defense pathways in *Arabidopsis thaliana*. Molecular Plant-Microbe Interactions 21, 208–218. ISSN: 0894-0282 (Jan. 2008).

33. Hartmann, A., Rothballer, M., Hense, B. A. & Schröder, P. Bacterial quorum sensing compounds are important modulators of microbe-plant interactions. English. Frontiers in Plant Science 5. ISSN: 1664-462X. doi:10.3389/fpls.2014.00131. https://www.frontiersin.org/articles/10.3389/fpls.2014.00131/full (2019) (2014).

34. Friesen, M. L. et al. Microbially mediated plant functional traits. Annual Review of Ecology, Evolution, and Systematics 42, 23–46 (2011).

35. Doty, S. L. en. in Endophytes of Forest Trees: Biology and Applications (eds Pirttilä, A. M. & Frank, A. C.) 151–156 (Springer Netherlands, Dordrecht, 2011). ISBN: 978-94007-1599-8. doi:10.1007/978-94-007-1599-8_9. https://doi.org/10.1007/978-94-007-1599-8_9 (2019).

36. Khan, Z. et al. Growth enhancement and drought tolerance of hybrid poplar upon inoculation with endophyte consortia. Current Plant Biology. Plant’s response to biotic and abiotic stresses and global climate change 6, 38–47. ISSN: 2214-6628 (Oct. 2016).

37. Kandel, S. L., Joubert, P. M. & Doty, S. L. Bacterial endophyte colonization and distribution within plants. en. Microorganisms 5, 77 (Dec. 2017).

38. Arnold, A. E. & Herre, E. A. Canopy cover and leaf age affect colonization by tropical fungal endophytes: ecological pattern and process in *Theobroma cacao* (Malvaceae). en. Mycologia 95, 388–398. ISSN: 0027-5514, 1557-2536 (May 2003).

39. Herre, E. A. et al. Ecological implications of anti-pathogen effects of tropical fungal endophytes and mycorrhizae. en. Ecology 88, 550–558. ISSN: 1939-9170 (2007).

40. Busby, P. E., Peay, K. G. & Newcombe, G. Common foliar fungi of *Populus trichocarpa* modify *Melampsora* rust disease severity. en. New Phytologist 209, 1681–1692. ISSN: 1469-8137 (Mar. 2016).

41. Busby, P. E. et al. Research priorities for harnessing plant microbiomes in sustainable agriculture. en. PLOS Biology 15, e2001793. ISSN: 1545-7885 (Mar. 2017).

42. Christian, N., Herre, E. A., Mejia, L. C. & Clay, K. Exposure to the leaf litter microbiome of healthy adults protects seedlings from pathogen damage. en. Proc. R. Soc. B 284, 20170641. ISSN: 0962-8452, 1471-2954 (July 2017).

43. Cheplick, G. P. & Cho, R. Interactive effects of fungal endophyte infection and host genotype on growth and storage in *Lolium perenne.* en. New Phytologist 158, 183–191. ISSN: 1469-8137 (2003).

44. Zahn, G. & Amend, A. S. Foliar fungi alter reproductive timing and allocation in *Arabidopsis* under normal and water-stressed conditions. en. bioRxiv, 519678 (Jan. 2019).

45. Christian, N., Herre, E. A. & Clay, K. Foliar endophytic fungi alter patterns of nitrogen uptake and distribution in *Theobroma cacao*. en. New Phytologist 222, 1573–1583. ISSN: 1469-8137 (2019).

46. Rosado, B. H. P., Almeida, L. C., Alves, L. F., Lambais, M. R. & Oliveira, R. S. The importance of phyllosphere on plant functional ecology: a phyllo trait manifesto. en. New Phytologist 219, 1145–1149. ISSN: 1469-8137 (2018).

47. Mejía, L. C. et al. Pervasive effects of a dominant foliar endophytic fungus on host genetic and phenotypic expression in a tropical tree. Frontiers in microbiology 5 (2014).

48. Dupont, P.-Y. et al. Fungal endophyte infection of ryegrass reprograms host metabolism and alters development. en. New Phytologist 208, 1227–1240. ISSN: 1469-8137 (Dec. 2015).

49. Dinkins, R. D., Nagabhyru, P., Graham, M. A., Boykin, D. & Schardl, C. L. Transcriptome response of *Lolium arundinaceum* to its fungal endophyte *Epichloë coenophiala*. en. New Phytologist 213, 324–337. ISSN: 1469-8137 (Jan. 2017).

50. Welsh, S. North American species of Astragalus Linnaeus (Leguminosae): a taxonomic revision. (Brigham Young University, 2007).

51. Knaus, B. J. Morphometric architecture of the most taxon-rich species in the U.S. flora: *Astragalus lentiginosus* (Fabaceae). en. American Journal of Botany 97, 1816–1826. ISSN: 0002-9122, 1537-2197 (Nov. 2010).

52. Baucom, D. L., Romero, M., Belfon, R. & Creamer, R. Two new species of *Undifilum,* fungal endophytes of *Astragalus* (locoweeds) in the United States. Botany 90, 866–875. ISSN: 1916-2790 (Aug. 2012).

53. Woudenberg, J. H. C., Groenewald, J. Z., Binder, M. & Crous, P. W. *Alternaria* redefined. eng. Studies in Mycology 75, 171–212. ISSN: 0166-0616 (June 2013).

54. Cook, D., Gardner, D. R., Martinez, A., Robles, C. A. & Pfister, J. A. Screening for swainsonine among South American *Astragalus* species. Toxicon 139, 54–57. ISSN: 0041-0101 (Dec. 2017).

55. Lu, H. et al. Endogenous fungi isolated from three locoweed species from rangeland in western China. en. African Journal of Microbiology Research 11, 155–170. ISSN: 1996-0808 (Feb. 2017).

56. Harrison, J. G., Parchman, T. L., Cook, D., Gardner, D. R. & Forister, M. L. A heritable symbiont and host-associated factors shape fungal endophyte communities across spatial scales. en. Journal of Ecology 106, 2274–2286. ISSN: 1365-2745 (Nov. 2018).

57. Molyneux, R. J. & James, L. F. Loco intoxication: indolizidine alkaloids of spotted locoweed *(Astragalus lentiginosus).* en. Science 216, 190–191. ISSN: 0036-8075, 1095-9203 (Apr. 1982).

58. Cook, D. et al. Swainsoninine concentrations and endophyte amounts of *Undifilum oxytropis* in different plant parts of *Oxytropis sericea*. eng. Journal of Chemical Ecology 35, 1272–1278. ISSN: 1573-1561 (Oct. 2009).

59. Shade, A. & Handelsman, J. Beyond the Venn diagram: the hunt for a core microbiome. en. Environmental Microbiology 14, 4–12. ISSN: 1462-2920 (Jan. 2012).

60. Gardner, D. R., Molyneux, R. J. & Ralphs, M. H. Analysis of swainsonine: extraction methods, detection, and measurement in populations of locoweeds *(Oxytropis* spp.) eng. Journal of Agricultural and Food Chemistry 49, 4573–4580. ISSN: 0021-8561 (Oct. 2001).

61. Ralphs, M. H. et al. Relationship between the endophyte *Embellisia* spp. and the toxic alkaloid swainsonine in major locoweed species (*Astragalus* and *Oxytropis*). eng. Journal of Chemical Ecology 34, 32–38. ISSN: 0098-0331 (Jan. 2008).

62. Grum, D. S. et al. Influence of seed endophyte amounts on swainsonine concentrations in *Astragalus* and *Oxytropis* locoweeds. Journal of Agricultural and Food Chemistry 60, 8083–8089. ISSN: 0021-8561 (Aug. 2012).

63. Cook, D. et al. A swainsonine survey of North American *Astragalus* and *Oxytropis* taxa implicated as locoweeds. Toxicon 118, 104–111. ISSN: 0041-0101 (Aug. 2016).

64. Grum, D. S. et al. Production of the alkaloid swainsonine by a fungal endophyte in the host *Swainsona canescens.* eng. Journal of Natural Products 76, 1984–1988. ISSN: 1520-6025 (Oct. 2013).

65. Cook, D., Gardner, D. R. & Pfister, J. A. Swainsonine-containing plants and their relationship to endophytic fungi. Journal of Agricultural and Food Chemistry 62, 7326–7334. ISSN: 0021-8561 (July 2014).

66. Panaccione, D. G., Beaulieu, W. T. & Cook, D. Bioactive alkaloids in vertically transmitted fungal endophytes. en. Functional Ecology 28, 299–314. ISSN: 1365-2435 (Apr. 2014).

67. Thompson, D. C., Knight, J. L., Sterling, T. M. & Murray, L. W. Preference for specific varieties of woolly locoweed by a specialist weevil, *Cleonidius trivittatus* (Say). English. Southwestern Entomologist 20, 325–325 (1995).

68. Parker, J. E. Effects of insect herbivory by the four-lined locoweed weevil, Cleonidius trivittatus (say) (Coleoptera: Curculionidae), on the alkaloid swainsonine in locoweeds Astragalus mollissimus and Oxytropis sericea en. Google-Books-ID: ODORP-gAACAAJ. PhD thesis (New Mexico State University, Las Cruces, New Mexico, 2008).

69. Ralphs, M. H., Panter, K. E. & James, L. F. Feed preferences and habituation of sheep poisoned by locoweed. eng. Journal of Animal Science 68, 1354–1362. ISSN: 0021-8812 (May 1990).

70. Creamer, R. & Baucom, D. Fungal endophytes of locoweeds: a commensal relationship? Journal of Plant Physiology and Pathology 1. doi:http://dx.doi.org/10.4172/2329-955X.1000104 (2013).

71. Schulthess, F. M. & Faeth, S. H. Distribution, abundances, and associations of the endophytic fungal community of Arizona fescue (*Festuca arizonica*). Mycologia 90, 569–578. ISSN: 0027-5514 (July 1998).

72. Cook, D. et al. Effects of elevated CO2 on the swainsonine chemotypes of *Astragalus lentiginosus* and *Astragalus mollissimus*. en. Journal of Chemical Ecology 43, 307–316. ISSN: 1573-1561 (Mar. 2017).

73. Oldrup, E. et al. Localization of endophytic *Undifilum* fungi in locoweed seed and influence of environmental parameters on a locoweed in vitro culture system. Botany 88, 512–521. ISSN: 1916-2790 (May 2010).

74. Högberg, P. 15N natural abundance in soil–plant systems. The New Phytologist 137, 179–203. ISSN: 1469-8137, 0028-646X (1997).

75. Harrison, J. G. et al. Deconstruction of a plant-arthropod community reveals influential plant traits with nonlinear effects on arthropod assemblages. en. Functional Ecology 32, 1317–1328. ISSN: 1365-2435 (May 2018).

76. Wang, Y. & Qian, P.-Y. Conservative fragments in bacterial 16S rRNA genes and primer design for 16S ribosomal DNA amplicons in metagenomic studies. en. PLOS ONE 4, e7401. ISSN: 1932-6203 (Oct. 2009).

77. White, T. J., Bruns, T., Lee, S. & Taylor, J. Amplification and direct sequencing of fungal ribosomal RNA genes for phylogenetics. In M. A. Innis, D. H. Glefand, J. J. Sninsky, and T. J. White [eds.], PCR protocols: A guide to methods and applications. (Academic Press, London, UK, 1990).

78. Edgar, R. C. Search and clustering orders of magnitude faster than BLAST. Bioinformatics 26, 2460–2461. ISSN: 1367-4803 (2010).

79. Edgar, R. C. & Flyvbjerg, H. Error filtering, pair assembly and error correction for next-generation sequencing reads. Bioinformatics, btv401. ISSN: 1367-4803 (2015).

80. Edgar, R. C. UNOISE2: improved error-correction for Illumina 16S and ITS amplicon sequencing. en. bioRxiv, 081257 (Oct. 2016).

81. Callahan, B. J., McMurdie, P. J. & Holmes, S. P. Exact sequence variants should replace operational taxonomic units in marker-gene data analysis. en. The ISME Journal 11, 2639. ISSN: 1751-7362 (July 2017).

82. Edgar, R. SINTAX: a simple non-Bayesian taxonomy classifier for 16S and ITS sequences. en. bioRxiv, 074161 (Sept. 2016).

83. Nilsson, R. H. et al. The UNITE database for molecular identification of fungi: handling dark taxa and parallel taxonomic classifications. en. Nucleic Acids Research. doi:10.1093/nar/gky1022. https://academic.oup.com/nar/advance-article/doi/10.1093/nar/gky1022/5146189 (2018) (2018).

84. Deshpande, V. et al. Fungal identification using a Bayesian classifier and the Warcup training set of internal transcribed spacer sequences. en. Mycologia 108, 1–5. ISSN: 0027-5514, 1557-2536 (Jan. 2016).

85. DeSantis, T. Z. et al. Greengenes, a chimera-checked 16S rRNA gene database and workbench compatible with ARB. en. Applied and Environmental Microbiology 72, 5069–5072. ISSN: 0099-2240, 1098-5336 (July 2006).

86. Cole, J. R. et al. Ribosomal Database Project: data and tools for high throughput rRNA analysis. en. Nucleic Acids Research 42, D633 (Jan. 2014).

87. Machida, R. J., Leray, M., Ho, S.-L. & Knowlton, N. Metazoan mitochondrial gene sequence reference datasets for taxonomic assignment of environmental samples. en. Scientific Data 4, 170027. ISSN: 2052-4463 (Mar. 2017).

88. Ntertilis, M., Kirmitzoglou, I., Promponas, V. J., Kouvelis, V. & Typas, M. MitoFun: a curated resource of complete fungal mitochondrial genomes. (Web page: *http://mitofun.biol.uoa.gr/*) 2013. http://mitofun.biol.uoa.gr/.

89. Tatusova, T., Ciufo, S., Fedorov, B., O’Neill, K. & Tolstoy, I. RefSeq microbial genomes database: new representation and annotation strategy. en. Nucleic Acids Research 42, D553–D559. ISSN: 0305-1048, 1362-4962 (Jan. 2014).

90. Fordyce, J. A., Gompert, Z., Forister, M. L. & Nice, C. C. A hierarchical Bayesian approach to ecological count data: a flexible tool for ecologists. en. PLOS ONE 6, e26785. ISSN: 1932-6203 (Nov. 2011).

91. R Core Team. R: A language and environment for statistical computing tex.organization: R Foundation for Statistical Computing. https://www.R-project.org/ (Vienna, Austria, 2019).

92. Plummer, M. JAGS: A program for analysis of Bayesian graphical models using Gibbs sampling. en. Proceedings of the 3rd international workshop on distributed statistical computing 124. Citation Key Alias: plummer03, 1–8 (2003).

93. Plummer, M. rjags: bayesian graphical models using MCMC. R package version 3-15. *https://CRAN.R-project.org/package=rjags* 2015.

94. Geweke, J. Evaluating the accuracy of sampling-based approaches to the calculation of posterior moments (Federal Reserve Bank of Minneapolis, Research Department, Minneapolis, MN, USA, 1991).

95. Gelman, A. & Rubin, D. B. Inference from iterative simulation using multiple sequences. Statistical Science 7, 457–472. ISSN: 0883-4237 (1992).

96. Gloor, G. B. & Reid, G. Compositional analysis: a valid approach to analyze microbiome high-throughput sequencing data. Canadian Journal of Microbiology 62, 692–703. ISSN: 0008-4166 (Apr. 2016).

97. Jost, L. Entropy and diversity. Oikos 113, 363–375. ISSN: 1600-0706 (2006).

98. Marion, Z. H., Fordyce, J. A. & Fitzpatrick, B. M. A hierarchical Bayesian model to incorporate uncertainty into methods for diversity partitioning. en. Ecology 99, 947–956. ISSN: 1939-9170 (2018).

99. Harrison, J. G. et al. The many dimensions of diet breadth: phytochemical, genetic, behavioral, and physiological perspectives on the interaction between a native herbivore and an exotic host. PLOS ONE 11, e0147971. ISSN: 1932-6203 (Feb. 2016).

100. Breiman, L. Random forests. en. Machine Learning 45, 5–32. ISSN: 0885-6125, 1573-0565 (Oct. 2001).

101. Liaw, A. & Wiener, M. Classification and regression by randomForest. R news 2, 18–22 (2002).

102. Pearson, K. Mathematical contributions to the theory of evolution—on a form of spurious correlation which may arise when indices are used in the measurement of organs. en. Proceedings of the Royal Society of London 60. classic, 489–498. ISSN: 0370-1662, (Jan. 1897).

103. Fernandes, A. D. et al. Unifying the analysis of high-throughput sequencing datasets: characterizing RNA-seq, 16S rRNA gene sequencing and selective growth experiments by compositional data analysis. Microbiome 2, 15. ISSN: 2049-2618 (May 2014).

104. Cook, D. et al. Swainsonine and endophyte relationships in *Astragalus mollissimus* and *Astragalus lentiginosus*. Journal of Agricultural and Food Chemistry 59, 1281–1287. ISSN: 0021-8561 (Feb. 2011).

105. Ralphs, M. H., Cook, D., Gardner, D. R. & Grum, D. S. Transmission of the locoweed endophyte to the next generation of plants. Fungal Ecology 4, 251–255. ISSN: 1754-5048 (Aug. 2011).

106. Marion, Z. H., Fordyce, J. A. & Fitzpatrick, B. M. Extending the concept of diversity partitioning to characterize phenotypic complexity. The American Naturalist 186, 348–361. ISSN: 0003-0147 (2015).

107. Palti, J. Biological characteristics, distribution and control of *Leveillula taurica* (Lév.) Arn. Phytopathologia Mediterranea 10, 139–153. ISSN: 0031-9465 (1971).

108. Correll, J. C., Gordon, T. R. & Elliott, V. J. Host range, specificity, and biometrical measurements of *Leveillula taurica* in California. Plant Disease 71, 248–251 (1987).

109. Vincent, J., Weiblen, G. & May, G. Host associations and beta diversity of fungal endophyte communities in New Guinea rainforest trees. Molecular ecology. ISSN: 1365294X (2015).

110. Coleman-Derr, D. et al. Plant compartment and biogeography affect microbiome composition in cultivated and native *Agave* species. en. New Phytologist 209, 798–811. ISSN: 1469-8137 (2016).

111. Griffin, E. A. et al. Foliar bacteria and soil fertility mediate seedling performance: a new and cryptic dimension of niche differentiation. en. Ecology 97, 2998–3008. ISSN: 1939-9170 (2016).

112. Laforest-Lapointe, I., Paquette, A., Messier, C. & Kembel, S. W. Leaf bacterial diversity mediates plant diversity and ecosystem function relationships. en. Nature 546, 145–147. ISSN: 1476-4687 (June 2017).

113. Strong, D. R., Lawton, J. H. & Southwood, S. R. Insects on plants. Community patterns and mechanisms. (Blackwell Scientific Publicatons, Oxford, UK, 1984).

114. Carmona, D., Lajeunesse, M. J. & Johnson, M. T. Plant traits that predict resistance to herbivores. en. Functional Ecology 25, 358–367. ISSN: 1365-2435 (Apr. 2011).

115. Berry, D. & Widder, S. Deciphering microbial interactions and detecting keystone species with co-occurrence networks. English. Frontiers in Microbiology 5. ISSN: 1664302X. doi:10.3389/fmicb.2014.00219. https://www.frontiersin.org/articles/10.3389/fmicb.2014.00219/full (2019) (2014).

116. Trosvik, P. & de Muinck, E. J. Ecology of bacteria in the human gastrointestinal tract—identification of keystone and foundation taxa. Microbiome 3, 44. ISSN: 20492618 (Oct. 2015).

117. Banerjee, S., Schlaeppi, K. & Heijden, M. G. A. v. d. Keystone taxa as drivers of microbiome structure and functioning. En. Nature Reviews Microbiology 16, 567. ISSN: 1740-1534 (Sept. 2018).

118. Braun, K., Romero, J., Liddell, C. & Creamer, R. Production of swainsonine by fungal endophytes of locoweed. Mycological Research 107, 980–988. ISSN: 0953-7562 (Aug. 2003).

119. Noor, A. I., Nava, A., Cooke, P., Cook, D. & Creamer, R. Evidence for nonpathogenic relationships of *Alternaria* section *Undifilum* endophytes within three host locoweed plant species. Botany 96, 187–200. ISSN: 1916-2790 (Jan. 2018).

120. Kulpa, S. M. & Leger, E. A. Strong natural selection during plant restoration favors an unexpected suite of plant traits. en. Evolutionary Applications 6, 510–523. ISSN: 1752-4571 (2013).

121. Leger, E. A. & Baughman, O. W. What seeds to plant in the Great Basin? comparing traits prioritized in native plant cultivars and releases with those that promote survival in the field. Natural Areas Journal 35, 54–68. ISSN: 0885-8608, 2162-4399 (Jan. 2015).

122. Klypina, N., Pinch, M., Schutte, B. J., Maruthavanan, J. & Sterling, T. M. Water-deficit stress tolerance differs between two locoweed genera (*Astragalus* and *Oxytropis*) with fungal endophytes. Weed Science, 1–13. ISSN: 1550-2759 (June 2017).

123. Coley, P. D., Bryant, J. P. & Chapin, F. S. Resource availability and plant antiherbivore defense. en. Science 230, 895–899. ISSN: 0036-8075, 1095-9203 (Nov. 1985).

124. Stamp, N. Out of the quagmire of plant defense hypotheses. The Quarterly Review of Biology 78, 23–55. ISSN: 0033-5770 (Mar. 2003).

125. Yan, J. F., Broughton, S. J., Yang, S. L. & Gange, A. C. Do endophytic fungi grow through their hosts systemically? Fungal Ecology 13, 53–59. ISSN: 1754-5048 (2015).

126. Co, A. D., Vliet, S. v., Kiviet, D. J., Schlegel, S. & Ackermann, M. Short-range interactions govern cellular dynamics in microbial multi-genotype systems. en. bioRxiv, 530584 (Jan. 2019).

127. Schmid, J. et al. Host tissue environment directs activities of an *Epichloë* endophyte, while it induces systemic hormone and defense responses in its native perennial ryegrass host. Molecular Plant-Microbe Interactions 30, 138–149. ISSN: 0894-0282 (Dec. 2016).

128. Zamioudis, C. & Pieterse, C. M. J. Modulation of host immunity by beneficial microbes. Molecular Plant-Microbe Interactions 25, 139–150. ISSN: 0894-0282 (Oct. 2011).

129. Chisholm, S. T., Coaker, G., Day, B. & Staskawicz, B. J. Host-microbe interactions: shaping the evolution of the plant immune response. Cell 124, 803–814. ISSN: 00928674 (Feb. 2006).

130. Dodds, P. N. & Rathjen, J. P. Plant immunity: towards an integrated view of plant–pathogen interactions. en. Nature Reviews Genetics 11, 539–548. ISSN: 1471-0064 (Aug. 2010).

131. Nissinen, R., Helander, M., Kumar, M. & Saikkonen, K. Heritable *Epichloë* symbiosis shapes fungal but not bacterial communities of plant leaves. En. Scientific Reports 9, 5253. ISSN: 2045-2322 (Mar. 2019).

132. Kannadan, S. & Rudgers, J. A. Endophyte symbiosis benefits a rare grass under low water availability. en. Functional Ecology 22, 706–713. ISSN: 1365-2435 (2008).

133. Galperin, M., Graf, S. & Kenigsbuch, D. Seed treatment prevents vertical transmission of *Fusarium moniliforme*, making a significant contribution to disease control. en. Phytoparasitica 31, 344–352. ISSN: 1876-7184 (Aug. 2003).

134. Barillas, J. R. V., Paschke, M. W., Ralphs, M. H. & Child, R. D. White locoweed toxicity is facilitated by a fungal endophyte and nitrogen-fixing bacteria. en. Ecology 88, 1850–1856. ISSN: 1939-9170 (July 2007).

